# Lipids Maintain Genomic Stability and Developmental Potency of Murine Pluripotent Stem Cells

**DOI:** 10.1101/2022.08.12.503780

**Authors:** Liangwen Zhong, Miriam Gordillo, Xingyi Wang, Yiren Qin, Yuanyuan Huang, Alexey Soshnev, Ritu Kumar, Gouri Nanjangud, Daylon James, C. David Allis, Todd Evans, Bryce Carey, Duancheng Wen

**Author notes:** These authors contributed equally (Miriam Gordillo, Xingyi Wang, Yiren Qin). Correspondence should be addressed to: Duancheng Wen, Ph.D.,; Bryce Carey, Ph.D., Todd Evans, Ph.D.

## Abstract

Lipids play vital roles in cellular homeostasis and regulate pluripotency of human stem cells. However, the impact of lipids on murine pluripotent stem cells is unclear. While Mek1/2 and Gsk3β inhibition (“2i”) supports the maintenance of murine embryonic stem cells (ESCs) in a homogenous naïve state, prolonged culture in 2i results in aneuploidy and DNA hypomethylation that impairs developmental potential. Additionally, 2i fails to support derivation and culture of fully potent female ESCs. Here we find that mouse ESCs cultured in 2i/LIF supplemented with lipid-rich albumin (AlbuMAX) undergo pluripotency transition yet maintain genomic stability and full potency over long-term culture. Mechanistically, lipids in AlbuMAX impact intracellular metabolism including nucleotide biosynthesis, lipid biogenesis, and TCA cycle intermediates, with enhanced expression of ZCAN4 and DNMT3s that prevent telomere shortening and DNA hypomethylation. In concert with 2i, lipids induce a formative-like pluripotent state through direct stimulation of Mek-mediated Erk2 phosphorylation, which also alleviates X chromosome loss in female ESCs. Importantly, both male and female “all-ESC” mice can be generated from *de novo* derived ESCs using AlbuMAX-based media. Our findings underscore the importance of lipids to pluripotency and link nutrient cues to genome integrity in early development.

## INTRODUCTION

Lipids play vital roles in the maintenance of cellular homeostasis by serving as an energy source through mitochondrial fatty acid oxidation, enhancing intracellular signal transduction, and providing macromolecules for membrane biosynthesis during growth and proliferation [1]. Lipids also affect stem cell pluripotency as demonstrated in recent reports that lipid supplementation in E8 medium induces differentiation of intermediate human ESCs into a primed stage [2, 3] and regulate human ESCs self-renewal [4]. However, the role of lipids on murine pluripotent stem cells is still poorly understood.

Pluripotency is the ability of a single cell to generate all cell lineages of an adult animal. Pluripotent murine ESCs were initially derived in media supplemented with fetal bovine serum (FBS) on feeder layers of mouse embryonic fibroblasts [5, 6]. Yet undefined serum components results in heterogeneous cultures and gradual loss of pluripotency [7]. The breakthrough discovery that inhibition of Mek1/2 and Gsk3β (2i) maintains murine ESCs in a more homogenous state of naive pluripotency [8], enabled stabilization and expansion of ground state ESCs *in vitro.* Recently, pluripotency has been described as a developmental continuum including three distinct phases: naïve, formative, and primed states, which correlate with murine embryonic stem cells (mESCs), formative pluripotent stem cells (fPSCs), and epiblast stem cells (EpiSCs), respectively [7–14]. Murine ESCs derived from the inner cell mass of E3.5-E4.5 blastocysts have unlimited self-renewal and differentiation capacity [5, 6, 8], as evidenced by their ability to generate all-ESC mice in tetraploid complementation assays [15–18]. Murine EpiSCs derived from post implantation epiblasts (E5.5–E8.25) are lineage primed for differentiation; their developmental potential is limited as they are unable to colonize blastocyst embryos [13, 14, 19, 20]. The recently proposed fPSCs are derived from E5.5-E6.5 embryos and are thought to possess unrestricted developmental potential. However, this has not been formally demonstrated, as the cells cannot be transitioned to a naïve state and they are less efficient compared to naïve cells for chimera contribution [9, 21, 22]. In these reports the fPSCs were derived by manipulating Activin A and Wnt signaling pathways, which may have different response thresholds for different PSC lines.

Defined serum-free 2i/LIF medium, which has been widely used in stem cell research, enables derivation of ESC lines from non-permissive strains that cannot be generated using serum-containing media [23–27]. Recently, however, two groups reported that prolonged culture of murine ESCs in 2i/LIF medium results in aneuploidy, loss of DNA methylation, and impaired developmental potential; moreover, the 2i/LIF system does not support the derivation of fully potent female ESCs [28, 29], suggesting a new approach is needed for deriving and maintaining stable ESCs of both sexes with full developmental potential.

Here, we provide definitive evidence that lipid supplementation (AlbuMAX, AX) maintains genomic stability and developmental potency over long term culture, and drives pluripotency transition through activated Erk signaling, which also supports female ESC X chromosome stability. Importantly, AX-induced formative-like ESCs are reversible by manipulating Fgf/Erk signaling, and AX-based medium is beneficial for both male and female ESC derivation, for culture and maintenance of full developmental potency in outbred or inbred strains.

## RESULTS

### AX improves developmental potential of murine ESCs

Murine naïve stem cells are maintained in a serum-free medium supplemented with GSK3β and Mek inhibitors (PD184352 and CHIR99021, 2i) and leukemia inhibitory factor (LIF) [8] (hereafter 2iL for the 2i/LIF medium or 2i-ESCs for the ESCs cultured in 2i/LIF medium). To test how lipids can influence pluripotent stem cells in serum-free media conditions, a chromatographically purified lipid-rich bovine serum albumin, AlbuMAX (A or AX), was added to 2iL culture medium (hereafter 2iLA for 2i/LIF+ AlbuMAX medium or AX-ESCs for ESCs cultured in 2iLA medium) (**Fig. 1a**). Male ESCs cultured in 2iLA medium maintained bright but flatter/larger colonies for over 15 passages, whereas the colonies in 2iL medium without AX acquired a flat and dark appearance (**Fig. 1b** **and Extended data Fig. 1a**). ESCs cultured in 2iLA proliferated more rapidly than in 2iL with doubling time at early passages (p3-p5) of 6.7h compared to 10.2h, respectively (**Extended data Fig. 1a**). Over continued passaging, the proliferation of AX-ESCs gradually slowed with doubling time increasing to 8.1h at passage 15 and reaching to 10.7h at passage 20 (**Extended data Fig. 1a**).

**Fig. 1.**
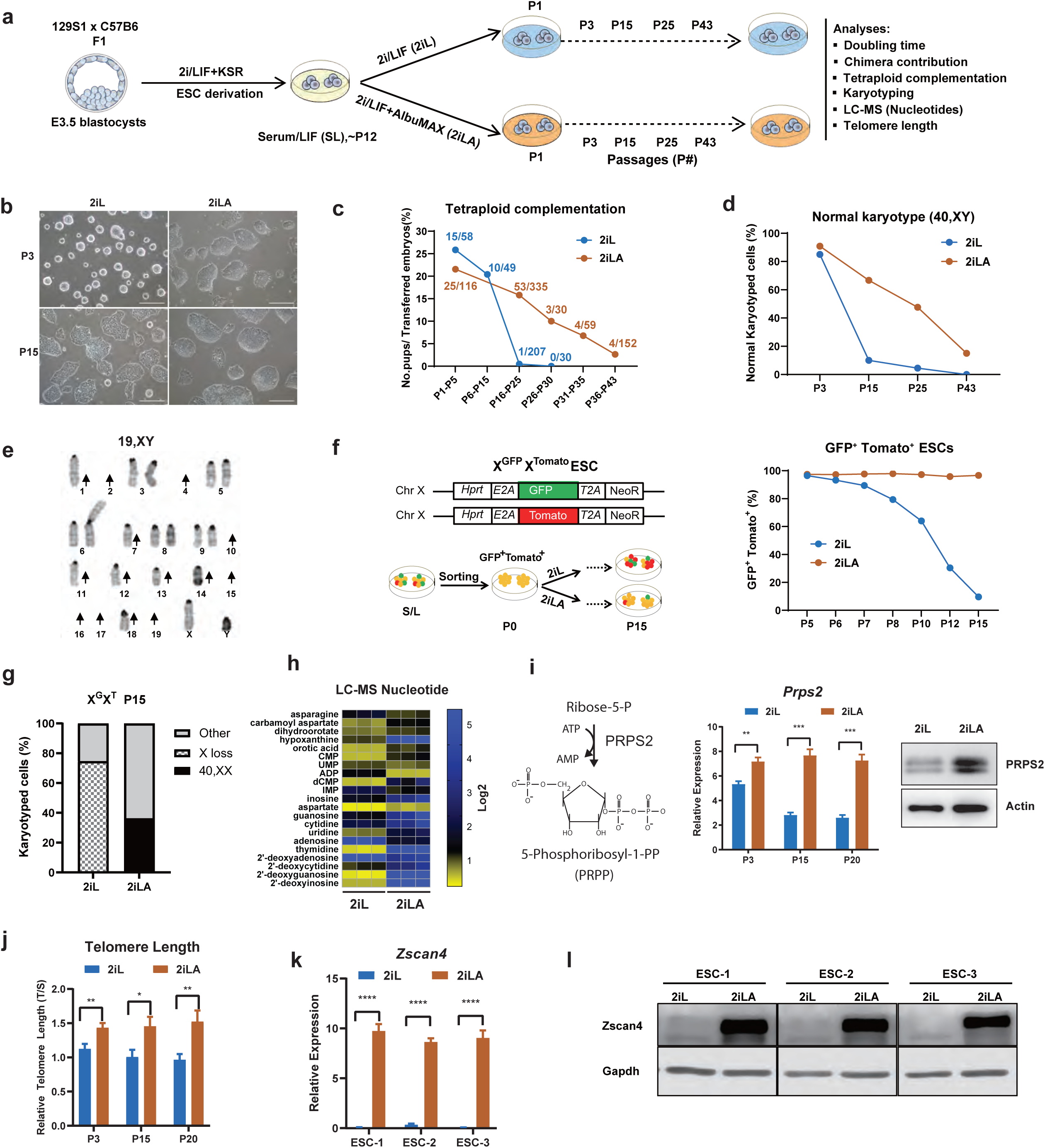
AX improves developmental potential and genomic stability of murine ESCs. **a,** Schematic illustration of the experimental design. **b,** Colony morphology for ESCs cultured in 2iL or 2iLA at passage 3 or passage 15. Scale, 25 µm. **c,** Developmental potential assessed by tetraploid complementation. The numbers at each timepoint represent total No. pups/No.embryos transferred. **d-e,** Analysis for normal karyotype (**d**) in 2iL and 2iLA at different passages (n=∼20 metaphases at each passage for 2iL or 2iLA). Representative severe chromosome loss (**e**) in 2iL at P43. Arrow: Chromosome loss. **f,** FACS analysis of XGXT reporter female ESC cultured in 2iL or 2iLA. The double reporter positive cells indicate both X chromosomes are maintained. **g,** Karyotyping of the XGXT ESCs cultured in 2iL and 2iLA at passage 15. **h,** LC-MS analysis of nucleotide pools for ESCs cultured in 2iL and 2iLA (P3). **i,** Prps2 expression levels by qRT-PCR assay or Western blot assay. **P<0.01, ***P<0.001, t-test. **j,** the telomere length analyzed by qRT-PCR relative to a single copy gene (T/S) for ESCs cultured in 2iL or 2iLA at different passages. * P<0.05, **P<0.01, t-test. **k-l,** Relative Zscan4 transcript (**k**) or protein (**l**) levels by qRT-PCR analysis or Western blot assay. ****P<0.0001, t-test.

After injection into blastocyst embryos, AX-ESCs gave rise to post-natal chimeras with germline transmission (**Extended data Fig. 1b).** To test for full developmental potency, AX-ESCs were assayed for the ability to generate all-ESC mice by tetraploid complementation. Both 2i-ESCs and AX-ESCs could generate all-ESC mice at high efficiency in early passages (p1–5) (**Fig.1c**). Consistent with previous studies [28, 29], prolonged culture of ESCs in 2iL demonstrated loss of developmental potential by 15 passages. Intriguingly, ESCs cultured in 2iLA medium retained the potential to generate all-ESC mice for over 20 passages (**Fig. 1c** **and Extended data Fig. 1c, 1d**). We frequently obtained as many as seven full-term live pups from a single surrogate mother (**Fig. 1c** **and Extended data Fig.1c),** efficiency similar to early passage 2i-ESCs. Although the potential for AX-ESCs to generate all-ESC mice decreased over time, it extended to passage 43 after 3 months of culture (**Fig. 1c** **and Extended data Fig. 1e)**.

### AX promotes karyotypic stability in ESC cultures

Because aneuploidy is the leading genetic cause of early pregnancy loss [30], we speculated that improved developmental potential of AX-ESCs may be due to improved genomic stability. We analyzed the karyotypes of ESCs cultured in 2iL and 2iLA media. Over 85% of ESCs cultured in either 2iL or 2iLA showed normal karyotype (40, XY) at passage 3 (P3), but by passage 15-25, nearly 95% of the cells maintained in 2iL exhibited aneuploidy (**Fig. 1d**), with trisomy 8 prevailing among aneuploid cells (**Extended data Fig. 2a-c**). By passage 43 nearly 100% of cells displayed aneuploidy, with a high prevalence of chromosome loss (**Fig. 1d-e, Extended data Fig. 2d-e**). Many of these ESCs had reduced chromosome content that approached haploid karyotype (**Fig. 1e** **and Extended data Fig. 2d-2e**). Remarkably, ESCs cultured in 2iLA medium had significantly higher number of cells exhibiting a normal karyotype, with over 68.3%, 50%, and 16.7% at passages 15, 25, and 43, respectively (**Fig. 1d**) indicating increased genome stability in cells grown in 2iLA. In addition, unlike 2i-ESCs, AX-ESCs maintained a near diploid karyotype (**Extended data Fig. 2d-2e)**. Female ESCs tend to quickly loss one of their two X chromosomes in 2iL medium [28, 29], so a dual reporter female ESC line (X^GFP^X^Tomato^, X^G^X^T^) [28] was employed to measure X chromosome loss by flow cytometry. Consistent with increased genomic instability, approximately 40% of cells at passage 10, and over 90% at passage 15, lost one X chromosome in ESCs cultured in 2iL medium (**Fig. 1f**). Notably, ESCs in 2iLA medium maintained two X chromosomes in over 95% of the cells with very little fluctuation from passage 3 to passage 15 (**Fig. 1f**). Karyotyping of the X^G^X^T^ ESCs at P15 confirmed the physical loss of X chromosome in 2i-ESCs and thus excluded the possibility of X chromosome inactivation in 2iL medium (**Fig. 1g**). Together, these data suggest that AX can efficiently prevent X chromosome loss in female ESCs and promotes karyotypic stability in ESC cultures.

### AX prevents nucleotide pool depletion and telomere shortening in 2iL medium

The reduced chromosome content of cultured 2i-ESCs (**Fig. 1e** **and Extended data Fig. 2d-2e**), indicates a possible deficiency in DNA replication and repair, which could be caused by imbalanced endogenous nucleotide pools [31–33]. We performed liquid chromatography-mass spectrometry (LC-MS) comparing ESCs cultured in 2iL or 2iLA conditions. Steady-state endogenous nucleotide pools are severely depleted in 2i-ESCs compared to AX-ESCs (**Fig. 1h**). Most of the nucleotides including thymidine, 2’-deoxyguanosine, 2’-deoxyinosine, and the precursor aspartate, are dramatically reduced in 2i-ESCs. Speculating that nucleotide biosynthesis pathways are impacted in 2iL medium, resulting in depletion of endogenous nucleotide pools, we analyzed the expression of Prps2, an ATP-dependent enzyme in the syntheses of purines and pyrimidines from ribose 5-phosphate [34, 35]. Real-time quantitative RT-PCR (qPCR) and western blotting analyses showed that Prps2 is expressed at significantly lower levels in 2i-ESCs compared to AX-ESCs (**Fig. 1i**), supporting the hypothesis that nucleotide biosynthesis is attenuated in 2iL medium and maintained by AX supplementation.

Decreasing dNTP (nucleotide) levels can lead to telomere shortening that also contributes to chromosomal instability [31, 36–38]. Therefore, we analyzed the telomere length of 2i-ESCs and AX-ESCs over passages. The telomere length of 2i-ESCs was shorter than that of AX-ESCs (**Fig. 1j, Extended data Fig.2f**), since the telomere length of AX-ESCs was maintained while that of 2i-ESCs became incrementally shortened from passage 0 to 6 (**Extended data Fig. 2g).** Additionally, Zscan4, a DNA-binding protein that functions for telomere elongation and maintenance in ESCs [39, 40], was expressed at significantly lower levels in 2i-ESCs compared to AX-ESCs (**Fig.1k, Extended data Fig. 2h**). Western blotting assays confirmed the Zscan4 protein was present in ESCs cultured in 2iLA but missing in 2iL (**Fig.1l**). The Zscan4-GFP and MERVL-tdTomato double reporter ESC line [41], was used to detect ESC transiently expressed early-embryonic transcripts, *Zscan4c* [39] and *MERVL* endogenous retrovirus [42]. This demonstrated that AX-ESCs exhibit a higher percentage of the Zscan4 positive or Zscan4/MERVL double positive cells (**Extended data Fig. 2i)**. Thus, suppression of Zscan4 expression in 2iL medium may account for the telomere shortening over multiple passages, and AX supplementation alleviates this suppression, thereby preventing telomere shortening during ESC *in vitro* culture.

### AX induces the transition of ESCs from naïve to formative-like pluripotency

While AX-ESCs retained the potential to generate all-ESC mice (**Fig. 1c**), we observed that colonies of these ESCs no longer maintained the typical uniform and rounded naïve colony morphology. Analyses of the core pluripotency genes, Pou5f1 (Oct3/4) and Sox2 by immunostaining, qPCR and western blotting assays revealed that these genes are expressed at similar levels independent of culture medium (**Fig. 2a-2b**, **Extended data Fig. 3a).** However, genes considered as naïve markers, such as Nanog, Rex1 (Zfp42), Klf2, Klf4, Prdm14, Stat3, Essrb, and Nr0b1 are all expressed at lower levels in AX-ESCs, more similar to ESCs in serum/LIF medium (**Fig. 2a-2c and Extended data Fig. 3a)**, indicating that AX-ESCs have lost naïve stem cell gene signatures and likely exited from the naïve pluripotent state. Since AX-ESCs can contribute to germline and maintain the potential to generate all-ESC mice by tetraploid complementation (**Fig. 1c**), they are not yet transformed to a primed state, but are likely maintained in a distinct naïve-to-primed intermediate state of pluripotency.

**Fig. 2.**
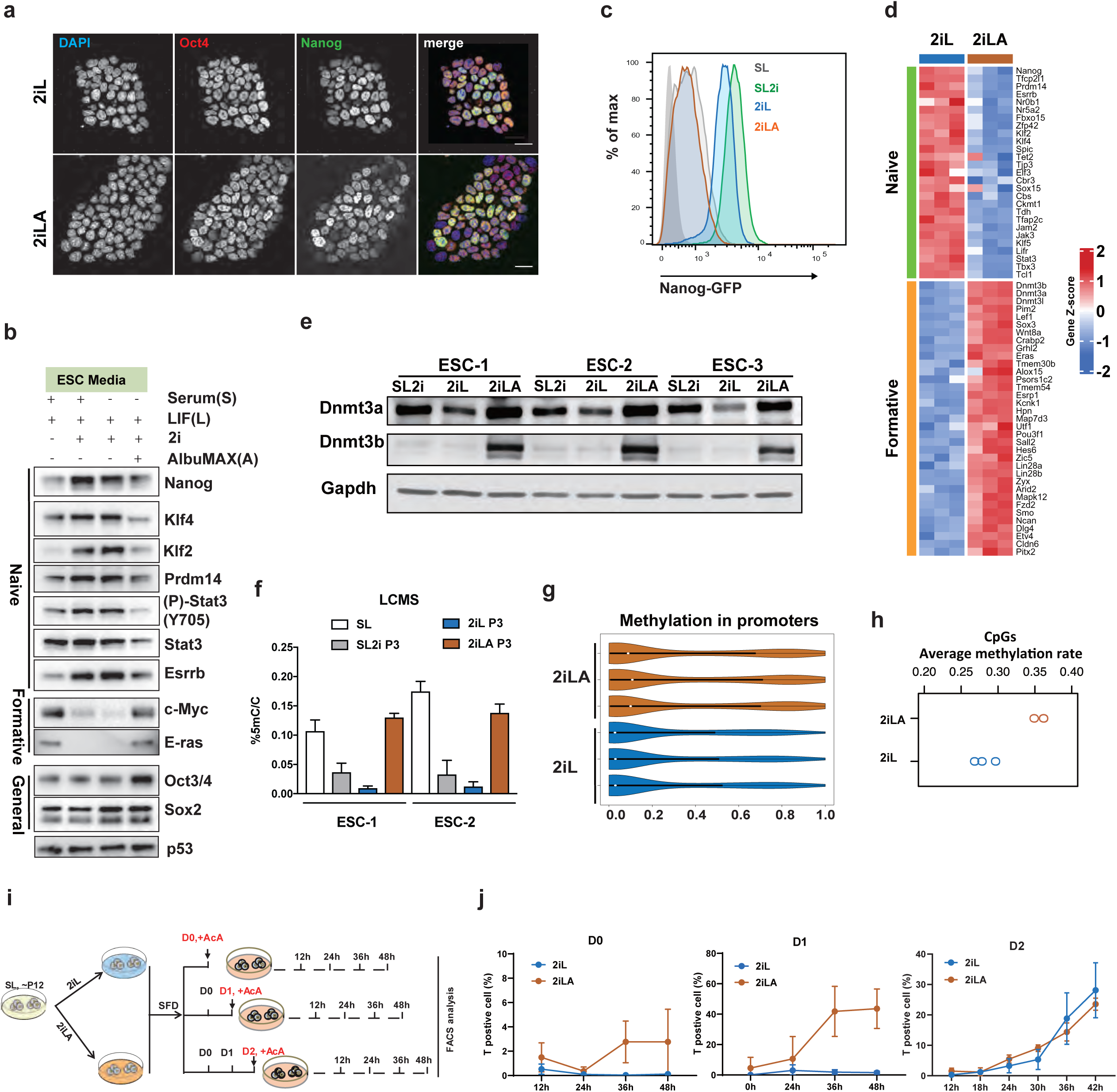
AX induces the transition of ESCs from naive to formative-like pluripotency. **a,** lmmunostaining for Oct4 (red) and Nanog (green) of ESCs cultured in 2il or 2iLA. DAPI (Blue) for cell nuclei. **b,** Western blotting of the naive,formative or general pluripotency marker protein level in SL,SL2i,2iL or 2iLA. **c,** FACS analysis using a GFP:Nanog ESC reporter line, the GFP fluorescence intensity is decreased in 2iLA. **d,** Heatmap of transcript levels from bulk RNA-seq for the naive and formative genes cultured in 2iL or 2iLA at P3. Three different ESC lines for biological replicates. **e,** Dnmt3a/b levels demonstrated by western blotting assays. **f,** LC-MS showing the global levels of 5mC/C ratio in genomic DNA from ESCs cultured in different media. **g,** Violin plots of the DNA methylation distribution rate at promoter regions in ESCs cultured in 2iL or 2iLA at P3 from reduced representation bisulfite sequencing (RRBS). Three biological replicates. **h,** RRBS for quantification of the average global methylation rate in all CpGs. Three biological replicates. **i,** Schematic of the assay to test responsiveness to Activin A for differentiation. **j,** FACS analysis of the mesendoderm differentiation marker brachyury (T) positive cells at different time points after Activin A (AcA) addition at Day0 (D0), Day1 (D1) or Day2 (D2) embryoid body formation. D0, D2 two biological replicates, D1 three biological replicates.

Formative pluripotency was recently proposed to describe an intermediate executive phase between the naïve and primed states [9–12, 21]. This phase of pluripotency is thought to be responsive to inductive cues for lineage differentiation and to express a group of unique formative genes [11, 43]. We performed RNA-seq of 2i-ESCs and AX-ESCs, finding 1637 genes that were differentially expressed and which are significantly enriched in pluripotency genes and the MAPK signaling pathway (**Extended data Fig. 3c).** Among these genes, 978 genes were upregulated in AX-ESCs, including most formative genes (**Fig. 2d**). Expression levels of formative genes such as Wnt8a, Pou3f1, Pim2, Sall2, Sox3, c-Myc, Eras, and Lef1, were all significantly higher in AX-ESCs, but silenced or expressed at low levels in 2i-ESCs (**Fig. 2b, 2d and Extended data Fig. 3b)**. Notably, genes expressed exclusively in primed stem cells, were not expressed in either 2i-ESCs or AX-ESCs (**Extended data Fig. 3d**).

Global increase of DNA methylation is anticipated in formative stem cells [11]. Expression levels for Dnmt3a, Dnmt3b and Dnmt3l are all higher in AX-ESCs compared to ESCs cultured otherwise, both in transcripts (**Extended data Fig.3b**) and protein (**Fig. 2e**). In agreement with these results, compared to the reduced total 5-methylcytosine (5mC) in 2iL ESCs, AX-ESCs exhibit a significant increase similar to levels seen with serum/LIF ESCs (**Fig. 2f**). Reduced-representation bisulfite sequencing (RRBS-Seq) confirmed the increase of global DNA methylation at promoters (**Fig. 2g**) and CpGs (**Fig. 2h**) in AX-ESCs. Of 5951 differentially methylated promoters, 5937 were hyper-methylated in AX-ESCs (>99%).

Responsiveness to inductive cues for lineage specification is another prominent feature that distinguishes naïve and formative stem cells [11]. To measure responsiveness, we adopted a standard directed differentiation protocol. ESCs were cultured in serum free differentiation (SFD) medium to form embryoid bodies (EBs), and Activin A was added at either day 0, day 1 or day 2 to induce differentiation toward mesendoderm, identified by Brachyury (T) expression (**Fig. 2i**). T positive cells were analyzed by flow cytometry to determine the differentiation efficiency. As expected, 2i-ESCs were not responsive to stimulus in the Day 0 group, failing to generate T positive cells (<0.1%) at the 4 timepoints (12h, 24h, 36h, 48h post Activin A induction) (**Fig. 2j**). Intriguingly, AX-ESCs were responsive to Activin A induction though not at high efficiency; approximately 5% T positive cells were found at 48h post induction (**Fig. 2j**). Notably, differentiation efficiency was remarkably increased for AX-ESCs when induced at Day 1 (after 24 hours in SFD medium), generating over 40% T positive cells at 36-48h post induction (**Fig. 2j**), whereas induction of the 2i-ESCs remained inefficient (∼5%) (**Fig. 2j**). The differentiation efficiency for 2i-ESCs significantly increased and reached comparable levels of AX-ESCs only when induced at Day2 (**Fig. 2j**), both showing similar differentiation kinetics and producing over 20% T positive cells 42h after induction (**Fig.2j**). These results are consistent with AX-ESCs acquiring a formative pluripotent state that ensures relatively rapid response to inductive cues and execution of lineage fate decisions. Taken together, the results indicate that AX-ESCs lose naïve stem cell identity and acquire a new formative-like intermediate state of pluripotency.

### Lipids in AX drive pluripotency transition

Previous studies of human pluripotent stem cells demonstrated that lipids in AX, not albumin (BSA), support self-renewal and facilitate pluripotent cell transitions [2, 4]. To test whether lipids are essential in the murine system, we first deproteinized AX (dep-AX) in a manner that can efficiently remove proteins including albumin from AX, as previously described [2]. ESCs cultured in 2iL medium supplemented with dep-AX retained a formative colony morphology (**Fig. 3a**), while ESCs cultured in 2iL supplemented with fatty acid free BSA (BSA) maintained a naïve colony morphology (**Fig.3a**). Addition of chemical defined lipid components (CDLC) plus BSA in 2iL medium, specifically Lipid Mixture 1 which contains non-animal derived fatty acids (arachidonic, linoleic, linolenic, myristic, oleic, palmitic, stearic, cholesterol, tocopherol acetate and Pluronic) could also induce the change of colony morphology (although not as dramatic as with AX), clearly distinct from naïve colony morphology, being flatter and much more irregularly shaped (**Fig. 3a**). Therefore, lipids, not albumin, induce the change of colony morphology for murine ESCs cultured in 2iLA medium.

**Fig. 3.**
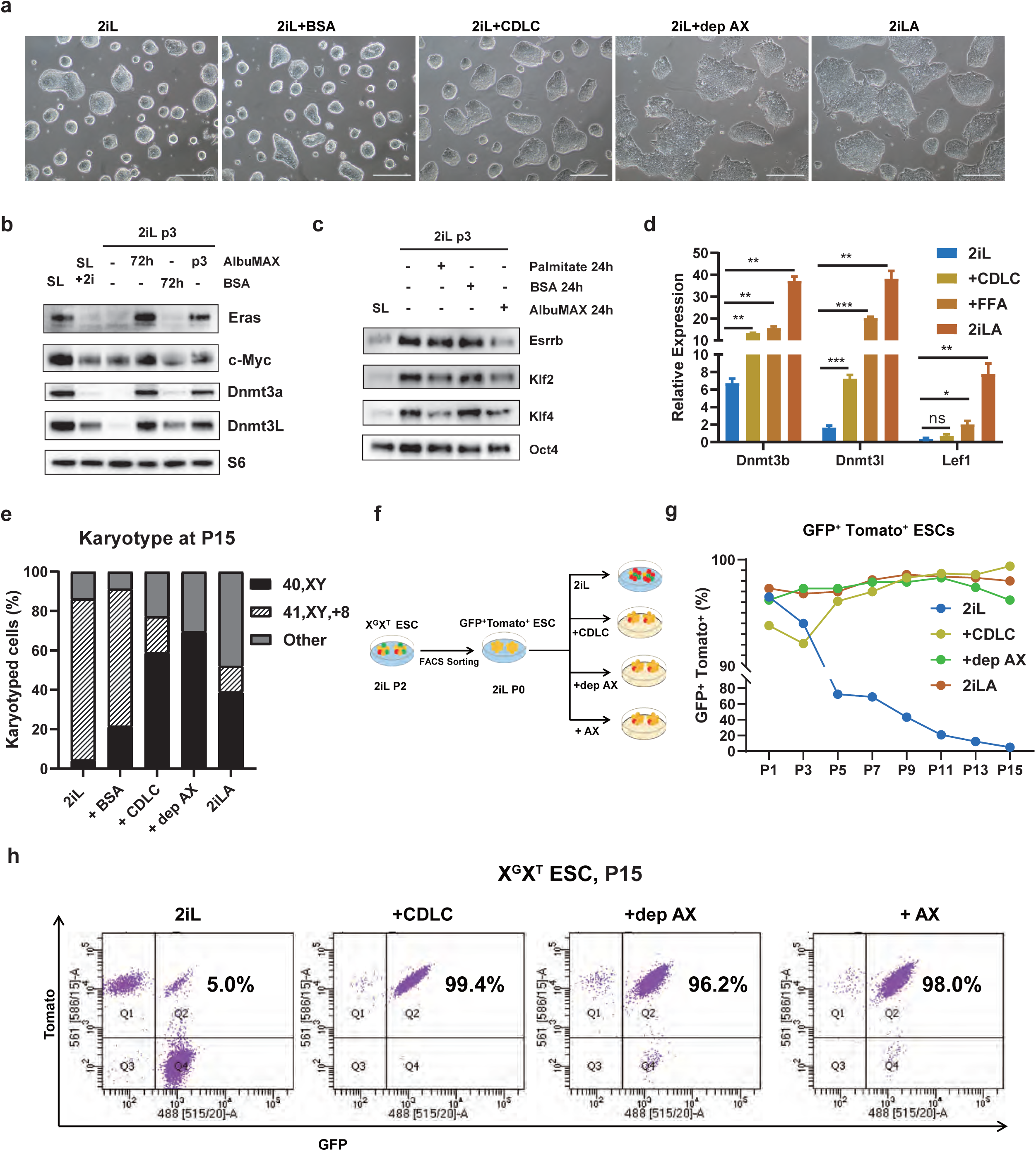
Lipids in AX drive the pluripotency transition and promote genomic stability. **a,** Colony morphology is shown for ESCs cultured in 2iL, 2iL+BSA (0.1%fatty acid free BSA), 2iL+CDLC (0.1%BSA+2% chemical defined lipid component), 2iL+dep-AX (deproteinized AX), and 2iLA. Scale bar, 25 µm. **b,** Western blotting assay to evaluate expression levels of Eras, c-Myc, Dnmt3a and Dnmt31. **c,** Western blotting assay to evaluate expression levels of Esrrb, Klf2, Kif 4 and Oct4. **d,** qRT-PCR expression levels of the formative marker genes *Dnmt3b*, *Dnmt3l*, and *Lef1* in ESCs cultured in 2iL, 2iL+CDLC, 2iL+dep-AX, or 2iLA. **e,** Karyotyping analyses of ESCs at P12 after culture in 2iL, 2iL+BSA (0.1%fatty acid free BSA), 2iL+CDLC (0.1%BSA +2%Chemical defined lipid component), 2iL+dep-AX (deproteinized AX), or 2iLA. **f,** Schematic of the experiments using X^G^X^T^ reporter ESCs to monitor the X chromosome loss. **g,** FACS analyses of X chromosome loss for X^G^X^T^ ESCs cultured in different media. **h,** Flow cytometry analysis of the X^G^X^T^ ESCs showing GFP+Tomato+ ESCs at passage 15.

We next analyzed the impact of lipid supplementation on expression of pluripotency genes. Western blot analysis showed that the protein levels of formative genes Eras, c-Myc, Dnmt3a, and Dnmt3L were increased (**Fig. 3b**) while those of naïve genes Esrrb, Klf2, and Klf4 were decreased in 2iL medium supplemented with AX or the saturated fatty acid palmitate (**Fig. 3c**); supplementing only BSA in 2iL medium had no effect on expression of these genes (**Fig. 3b, 3c**). qPCR assays demonstrated that levels of the formative genes *Dnmt3s* and *Lef1* were enhanced in 2iL medium supplemented with CDLC or free fatty acids (FFA) (**Fig. 3d**). Therefore, lipids induce the expression of formative genes and decrease expression of naïve genes, indicating that lipids are sufficient, independent of BSA, to induce the pluripotency transition to formative-like cells in 2iLA medium.

### Lipids in AX impact intracellular metabolism and enhanced non-canonical TCA cycle metabolism

In addition to increased steady-state levels of nucleotide pools, LC-MS analysis also showed that treatment with AX enhanced the steady-state levels of the non-canonical TCA cycle metabolites citrate and cis-aconitic acid (**Extended data Fig. 4a, b**), which were recently reported to be engaged in the exit of naïve pluripotency [44]. qRT-PCR analysis of TCA cycle enzymes showed that AX treatment did not alter expression of most genes with the exception of succinate dehydrogenase (*SDHB*), which was upregulated after just 24-hour treatment and sustained through passage 3 (**Extended data Fig 4c**). In serum-free 2iL medium cells must expend high levels of reducing equivalents in the form of NADPH to synthesize lipids de novo. Consistent with high levels of de novo lipogenesis, ACLY, FASN, SCD1, SCD2 protein levels are increased in 2iL while treatment with AX induced a strong downregulation indicating a reduction in capacity for lipid biosynthetic reactions (**Extended Fig 4d-e**). Moreover, two metabolic intermediates associated with lipid metabolism, glycerol-3-phosphate and phosphoethanolamine, were also reduced in AX treated cells (**Extended data Fig 4f**). LC-MS also revealed a steady state reduction in amino acid abundance (**Extended data Fig 4g**), potentially a result of increased anabolic reactions and protein synthesis in AX-ESCs expressing high levels of c-Myc and ERas.

### Lipids in AX promote genomic stability and maintain two active X chromosomes in female ESCs

We next evaluated the effect of lipids on genomic stability. Male ESCs expanded under 2iL (unsupplemented), AX, dep-AX or CDLC conditions were karyotyped at P12. As expected, the XY ESCs cultured in 2iL medium became aneuploid (95%), while 40% of cells in 2iLA medium maintained a normal karyotype (**Fig.3e**). Medium supplemented with CDLC or dep-AX promoted retention of normal ESC karyotype (60% and 70%, respectively) (**Fig. 3e**). These results demonstrated that both CDLC and dep-AX can significantly promote the karyotypic stability of male ESCs comparable to AX supplementation of 2iL medium.

To investigate the effect of lipids on X chromosome loss in female ESCs, we adopted the X^G^X^T^ double reporter system to monitor X chromosome loss during *in vitro* culture (**Fig. 1f**). The GFP and Tomato double positive cells were sorted at passage 2 in 2iL and expanded in 2iL for 2 passages. The cells were then split at the same density into 2iL medium alone, or supplemented with AX, dep-AX or CDLC (denoted as passage 0, P0), followed by FACS analyses to assess X chromosome loss over the course of *in vitro* expansion (**Fig. 3f**). The double positive cell population was incrementally depleted from P1 to P15 in 2iL medium, but it was maintained at a high percentage (over 95%) in 2iLA medium (**Fig. 3g**). Notably, both dep-AX and CDLC supplementation also preserved a high double positive cell population (over 95%) (**Fig. 3g**), showing that lipids efficiently prevent X chromosome loss in supplemented 2iL medium. Therefore, lipids can mitigate aneuploidy and stabilize the genome of murine ESCs cultured in 2iL medium, supporting the notion that lipids are responsible for the effects of AX on genomic stability.

### Lipid-induced formative-like pluripotency is maintained in concert with 2i

To explore how AX interacts with the two inhibitors (2i) to induce pluripotency transition and maintain genomic stability, we removed PD, CHIR or both from 2iLA medium (designated as CLA, PLA or LA). Withdrawal of either or both inhibitors from 2iL or 2iLA media led to loss of a typical formative colony morphology as seen in 2iLA (**Fig. 4a, Extended data Fig.5a**). However, LA medium supported proliferation and long-term maintenance of ESCs, while with only LIF in the basic media without AX massive cell death occurred during the first several passages (**Fig. 4a**). Expression levels of the formative marker genes *Dnmt3b*, *Lef1* and*Pou3f1* were significantly changed compared to AX-ESCs (**Extended data Fig. 5b**). The growth curve showed CLA, or PLA ESC number were significantly decreased, possible because of the cell death increase of the differentiated ESCs (**Fig. 4a-b**). Moreover, LA-ESCs responded more rapidly to Activin A stimulus than naïve 2i-ESCs, even faster than AX-ESCs (**Fig.4c**). Therefore, LA-ESCs lose naïve stem cell identity and transition to a pluripotency state similar to AX-ESCs.

**Fig. 4.**
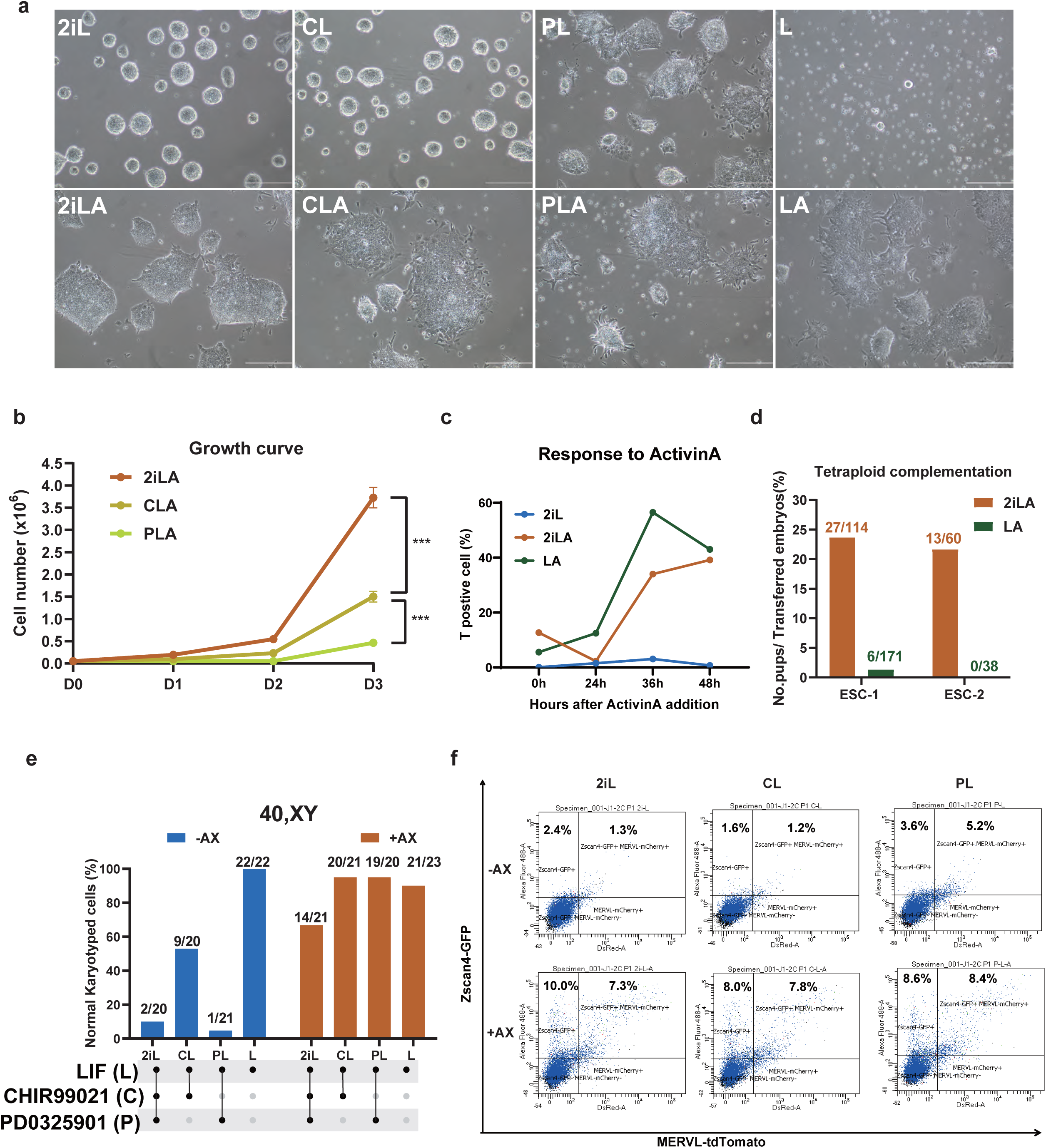
Lipid-induced formative pluripotency is maintained in concert with 2i. **a,** shown are representative colony morphologies of ESCs cultured in 2iL (CHIR+PD+LIF), CL (CHIR+LIF), PL (PD+LIF), L (LIF) with or without 1%AlbuMAX (A) at passage 3 or 4. Scale, 25 µm. **b,** the cell growth curves of ESCs cultured in 2iLA, CLA or PLA at passage 3. ***P<0.001, t-test. **c,** FACS analysis of the mesendoderm differentiation marker brachyury (T) positive cells at different time points after Activin A addition for ESCs cultured in 2iL, 2iLA or L to test the responsiveness to directed differentiation. **d,** Tetraploid complementation assay for two ESC lines cultured in 2iLA or LA. **e,** Karyotyping analyses of ESCs cultured in different media with AlbuMax (+AX) or without (-AX) after 15∼16 passages. Normal karyotype/Total analysis metaphases. n=∼20 metaphases for each condition. 2iL: CHIR+PD+LIF, CL: CHIR+LIF, PL: PD+LIF, L: LIF. **f,** FACS analysis to evaluate the Zscan4 positive cell percentage using the Zscan4-GFP and MERVL-tdTomato reporter ESC line cultured in different media at P1.

The developmental potential of LA-ESCs was assessed using tetraploid complementation. Live pups were obtained from the LA-ESCs when used at early passage (P3) in one cell line (ESC-1). However, this was very inefficient compared to using AX-ESCs, with 4n competence completely lost by passage 10 (**Fig. 4d**). Indeed, a second cell line failed to generate live pups even at early passages (P3-P6) (**Fig. 4d**). Karyotyping analyses of LA cultured ESCs revealed that the genome is stable - over 90% of cells maintained normal karyotype at P15 (**Fig. 4e**), suggesting that the loss of 4n competence in LA-ESCs is due to epigenetic changes rather than genetic deterioration. Therefore, ESCs cultured in LA medium maintain formative-like colony morphology and express formative-like marker genes, yet do not have the full potency as seen with ESCs cultured in 2iLA medium. This suggests that the formative pluripotency induced by lipids in 2iLA medium is maintained in concert with the two inhibitors of Mek/Erk and GSK3β signaling pathways. Notably, karyotypic analyses showed that addition of either PD or CHIR promotes abnormal karyotypes in 2iL medium, (especially PD). However, AX can blunt the adverse effects of PD and/or CHIR on genomic stability. Most of the L-ESCs or LA-ESCs maintained karyotype stability (**Fig. 4e, Extended data Fig.5c**). Additionally, AX supplement enhanced the Zscan4-GFP positive population (**Fig. 4f**) in Zscan4-GFP and MERVL-tdTomato double reporter ESCs (**Extended data Fig. 2i**) regardless of the inhibitors. The data indicate that the ability of lipids to increase ZSCAN4 expression levels and maintain genomic stability is independent of GSK3 and Erk signaling.

### Fatty acid oxidation (FAO) is dispensable for lipid-induced pluripotency transition

To determine whether lipids or lipid metabolism (lipid metabolites) drive pluripotency transition, we inhibited carnitine palmitoyltransferase (CPT1) with Etomoxir (ETO) (**Extended data Fig. 6a**), an irreversible inhibitor of CPT1 on the inner face of the outer mitochondrial membrane [45]. Pluripotency transition from naïve to formative state was observed in ESCs cultured in 2iLA medium supplemented with ETO (**Extended data Fig. 6b, 6c).** ACAA2 catalyzes the last step of the mitochondrial fatty acid beta oxidation spiral (**Extended data Fig. 6a**); inhibition of ACCA2 with Trimetazidine (TMZ) [46] also did not block the pluripotency transition in 2iLA medium (**Extended data Fig. 6b)**. To validate these results, we took a genetic approach by disruption in ESCs of the *Cpt1a* gene using CRISPR/Cas9 (**Extended data Fig. 6d)**. Consistent with inhibitor results, the *Cpt1a* null ESCs displayed a formative-like colony morphology in 2iLA medium (**Extended data Fig. 6e)** and increased expression levels of formative gene *Dnmt3b*, based on qRT-PCR (**Extended data Fig. 6f)**. Therefore, fatty acid oxidation is dispensable for the lipid-induced formative pluripotency, suggesting that lipids, rather than lipid metabolism, drive pluripotency transition.

### Lipids drive pluripotency transition via stimulating Mek-mediated Erk2 phosphorylation

While Mek/Erk signaling is thought to be inhibited by PD in 2iLA medium, western blotting assays indicated that phosphorylated-Erk1/2 (p-Erk1/2) was significantly increased in ESCs cultured in 2iLA medium compared to serum/LIF+2i medium (**Fig. 5a**). This suggests that AX blunts the effect of PD on the inhibition of MAPK signaling. The kinetics of AX and p-Erk1/2 signaling was explored by performing a time course experiment focused on genes that are upstream of Erk1/2. Mek1/2, p-Mek1/2, and p-Raf (Erk upstream genes), did not respond to AX addition at the protein level, while levels of p-Erk1/2 increased as early as 10 min and peaked from 20 min through 5h post AX supplementation (**Fig. 5b**), with higher p-Erk1/2 levels maintained in 2iLA medium thereafter (**Fig. 5b**). Notably, qPCR analyses showed that expression of *Dnmt3a/b/l* were not changed until 1h after AX addition (**Extended data Fig. 7a**). Therefore, the increase of p-Erk1/2 precedes the upregulation of formative genes, suggesting that the change of p-Erk1/2 levels is a direct response to AX addition, rather than a secondary effect of pluripotency transition.

**Fig. 5.**
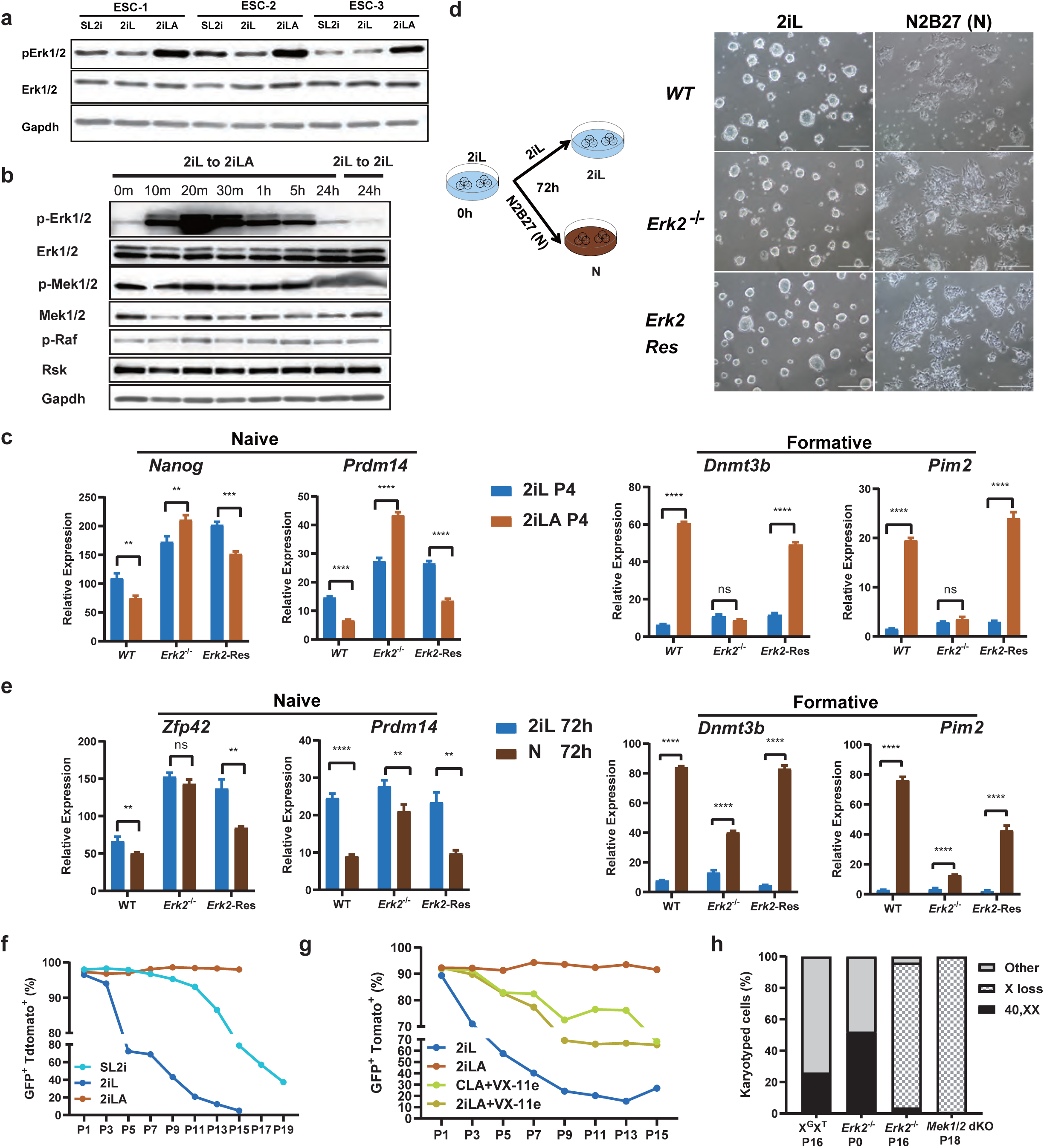
Mek/Erk signaling is essential for lipid induced formative transition. **a-b,** Western blotting for Erk1/2 and p-Erk1/2 proteins in ESCs cultured in 2iL or 2iLA at passage 3 (a) or for Mek/Erk signaling related proteins in ESCs after switching 2iL to 2iLA (b). m, minutes; h: hours. **c,** qRT-PCR for the naive marker genes *Nanog*, *Prdm14* and formative marker genes *Dnmt3b*, *Pim2* for *WT*, *Erk2^-/-^*, *Erk2-Res* ESCs at passage 4 cultured in 2iL or 2iLA. *Erk2-Res*: *Erk2*-rescued in the *Erk1/2* mutant ESCs (Lentiviral-based shRNA against *Erk1* in the *Erk2^-/-^* ESCs). n= 3 experiments. ns, no significant difference, *P<0.05, **P<0.01, ***P<0.001, ****P<0.0001, student t-test. **d-e,** Schematic illustration of the experimental design and ESC colony morphology change (d), and qRT-PCR for the naive marker genes *Zfp42*, *Prdm14*, and the formative marker genes *Dnmt3b*, *Pim2* for WT, and *Erk2* ^-/-^*Erk2-Res* (e) after 2i and LIF removal for 72 hours. n= 3 experiments. **f-g,** FACS analysis of X^G^X^T^ reporter female ESCs cultured in SL2i, 2iL or 2iLA from P1 to over P15 (f) or in 2iL, 2iLA, 2iLA + VX-11e (Erk2 inhibitor) or CLA + VX-11e from P1 to P15 (g), CLA: CHIR 99021 +LIF+ AlbuMAX. **h,** Karyotyping of the X^G^X^T^, *Erk2-/-*, *Mek1*^-/-^*Mek2*^-/-^ double knockout (*Mek1/2* dKO) PSCs cultured in 2iLA over passage 15. n=∼20 metaphases for each cell line.

To investigate if Erk1/2 signaling is involved in the pluripotency transition induced by lipids, we used an ESC line in which *Erk2* is constitutively deleted and *Erk1* is knocked down by shRNA (designated *Erk1/2* mutant) [47]. With the *Erk1/2* mutant, both Erk1 and Erk2 proteins are depleted, while Erk2 activity can be rescued by adding back the *Erk2* cDNA (Erk2-Rescue, *Erk2-Res*) (**Extended data Fig. 7b),** as confirmed by the expression of *Spry4*, a direct target of the Erk1/2 signaling pathway (**Extended data Fig. 7c)**. When culturing *Erk1/2* mutant ESCs in 2iLA medium, the expression levels for formative genes *Wnt8a and Lef1* were relatively unchanged compared to the 2iL controls (**Extended data Fig. 7d**), suggesting that pluripotency transition is likely blocked in 2iLA medium in these cells.

Erk1/2 signaling is essential for self-renewal and maintenance of ESCs, and double knockout of *Erk1* and *Erk2* is not tolerated *in vitro* [48]. To circumvent this problem, we analyzed single *Erk2* knockout, as *Erk2* is thought to be the key gene regulating pluripotency transition [49, 50]. Gene expression analysis showed that *Erk2* deletion is sufficient to block response to lipid induced pluripotency transition (**Extended data Fig. 7d**). Remarkably, the pluripotency transition was severely impaired in *Erk2^-/-^* ESCs induced by lipids even at passage 4, as indicated by the sustained expression of naïve genes such as *Nanog* and *Prdm14*, while the expression levels of formative genes *Dnmt3b* and *Pim2* remained largely unchanged (**Fig. 5c**). To test if this defect is specific to lipid induced transition, we induced the *Erk2^-/-^* ESC differentiation by removal of 2i and LIF from the basic N2B27 culture medium (**Fig. 5d**). Under these conditions, *Erk2^-/-^* cells initiated differentiation and formed differentiated colonies similar to WT ESCs at 72h (**Fig. 5d**). qRT-PCR analyses of the *Erk2^-/-^* cells showed that the expression levels of naïve genes *Prdm14* and *Zfp42* were downregulated while those for the formative genes *Dnmt3b* and *Pim2* were significantly upregulated (**Fig. 5e**). However, the transcriptional changes in *Erk2^-/-^* cells after switching to N2B27 medium were relatively modest compared to WT cells, suggesting a delay in the pluripotency transition, as previously described for multiple mutants in the Fgf/Erk pathway including *Erk1 and Erk2* single mutants [51]. Finally, we examined the pluripotency transition induced by lipids in PSCs with *a* double knockout of *Mek1/2,* the only kinases known to be responsible for Erk1/2 phosphorylation in PSCs (**Extended data Figs. 7e-7f**). Analysis of pErk1/2 confirmed that phosphorylation of Erk1/2 requires p-Mek1/2 in 2iLA medium (**Extended data Fig. 7h**). The pluripotency transition was significantly impaired in 2iLA medium as indicated by expression levels of naïve and formative markers (**Extended data** **Fig. 7g**). Furthermore, when switched from 2iL to 2iLA, deletion of *Mek1/2* or *Erk2* resulted in smaller colonies compared to *WT* or *Erk2-Re*s, with reduced proliferation than the control, based on the colony size (**Extended data Fig. 7i**). Also, both the naive gene *Nanog* and the formative gene *Dnmt3l* were deregulated after switching into 2iLA (**Extended data Fig. 7j**). This result suggests that stimulation of Erk2 by lipids requires Mek. Together, our results indicate that lipids can stimulate Mek-mediated Erk2 phosphorylation which is essential for lipid-induced pluripotency transition.

### Erk1/2 suppression leads to X chromosome loss in female ESCs

The cause of X chromosome loss when female ESCs are cultured in 2iL is unclear, although it has been suggested to be related to Wnt and Mapk signaling [28]. We showed that adding lipids in 2iL medium efficiently prevents X chromosome loss (**Fig. 3g**), while p-Erk1/2 level is increased in this condition (**Fig 5a**). We hypothesized that suppression of Erk1/2 signaling in 2iL medium is responsible for X chromosome loss in XX ESCs, whereas lipid-induced increase of Erk1/2 signaling attenuates X chromosome loss. Using the X^G^X^T^ reporter ESC line, FACS analyses showed that ESCs cultured in 2iL or serum/LIF/2i (SL2i) with lower p-Erk1/2 underwent significant X chromosome loss by passage 15, while in 2iLA, ESCs maintained over 95% of two active X chromosomes (**Fig. 5f**). To determine if Erk1/2 signaling is associated with X chromosome loss, we inhibited Erk2 activity using VX-11e, a small molecule inhibitor of Erk2 [52] in 2iLA or CLA medium (2iLA+VX-11e or CLA+ VX-11e). Indeed, significant X chromosome loss was observed (over 30% loss of X chromosome after P9) in ESCs cultured in 2iLA+VX-11e medium (**Fig. 5g**). The result supports the notion that suppression of Erk signaling is associated with X chromosome loss in XX ESCs cultured in 2iL medium. Moreover, genetic deletion of Mek1/2 or Erk2 in ESCs demonstrated that nearly 100% of ESCs lost one X chromosome by passage 16 or 18 even when cultured in 2iLA medium (**Fig. 5h**), suggesting that Mek/Erk signaling is critical for maintenance of two X chromosomes in ESCs. Collectively, our results suggest that prolonged suppression of Mek/Erk signaling is responsible for the X chromosome loss in ESCs cultured in 2iL medium and lipids prevent X chromosome loss through increase of p-Erk1/2.

### Lipid-induced formative-like pluripotency is reversible

We tested if the naïve to formative pluripotent state induced by lipids could be reversed upon withdrawal of lipids. Indeed, AX-ESCs that maintained stable formative features for >10 passages formed typical naïve colony morphology when switched to 2iL medium (designated R-ESCs) (**Fig. 6a**). Bulk RNA-seq on these cells showed that upon withdrawal, naïve markers were enhanced whereas formative markers were decreased (**Fig. 6b**). Like 2i-ESCs, the endogenous nucleotide pools were significantly depleted in R-ESCs (**Fig. 6c**). Western blotting assays confirmed that in R-ESCs, expression of p-Erk1/2, Dnmt3a/b and Zscan4 decreased to levels found in naïve 2i-ESCs (**Fig. 6d**). Using the *in vitro* system described above to test responsiveness to differentiation stimuli (**Fig. 2i**), Activin A induction of T positive cells was decreased for R-ESCs (**Fig. 6e**). Our results suggest that the formative state of AX-ESCs can revert fully back to a naïve state by withdrawal of AX from the culture medium.

**Fig. 6.**
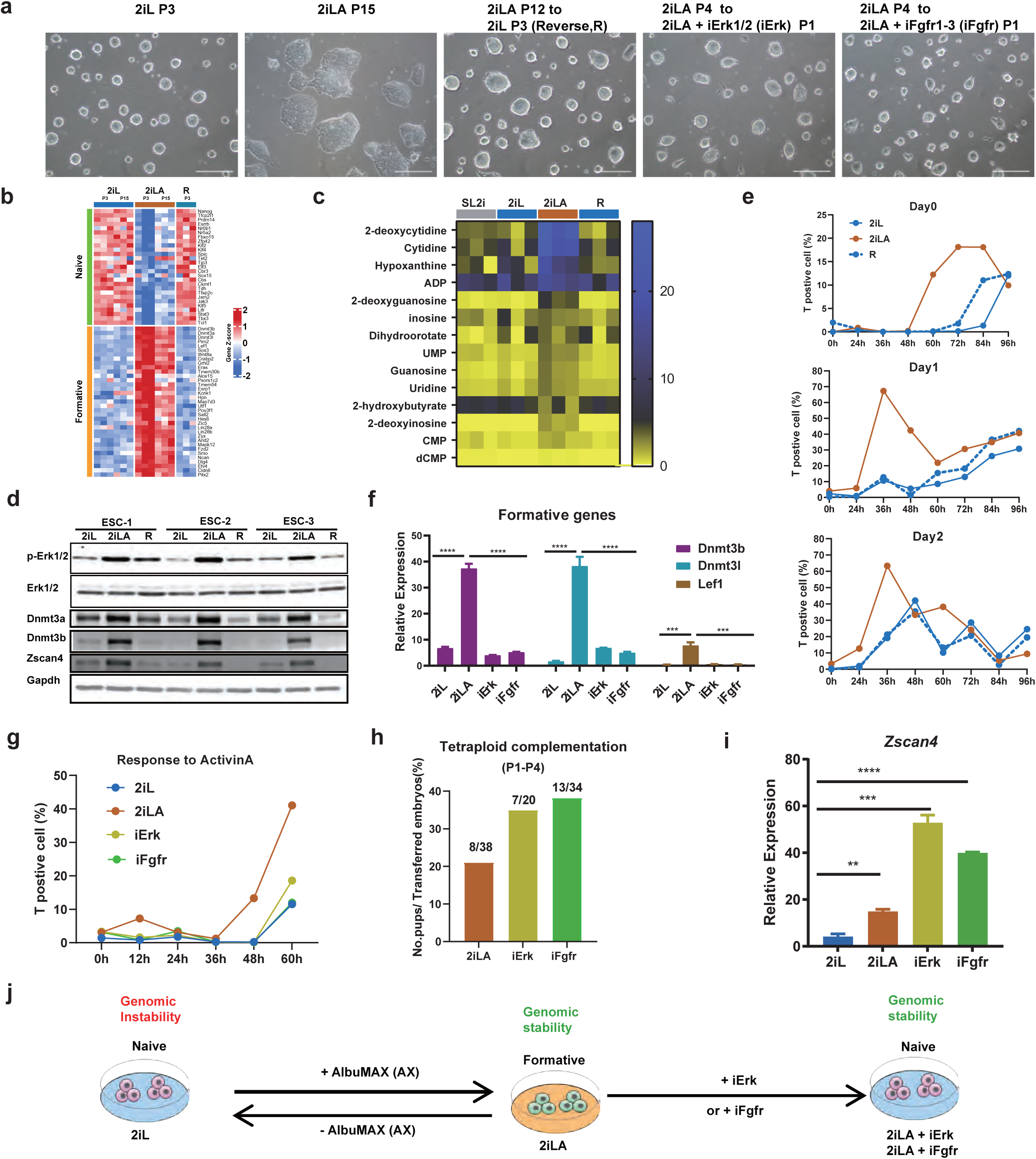
Lipid-induced pluripotency transition is restrained through manipulating Fgf/Erk signaling. **a,** The colony morphology of ESCs cultured in 2iL at P3, 2iLA at P15, reversing 2iLA at P12 to 2iL at P3 (R), 2iLA+iErk (1μM SCH772984, iErk) at P1 and 2iLA+iFgfr1-3(1μM AZD4547, iFgfr1-3) at P1. Scale, 25 μm. **b-c,** Heatmap for the naive and formative genes based on bulk RNA-seq (b) or nucleotides analysis by LC-MS. (c) for ESCs at P3 or P15 in 2iL, 2iLA or R-ESC (R, Reversing 2iLA at P12 to 2iL at P3). n= 3 biological replicates. **d,** Western blotting for proteins for ESCs in 2iL or 2iLA at P15 and R-ESCs at P3. **e,** FACS for the differentiation marker brachyury (T) positive cells after adding Activin A after Day0, Day1, or Day2 embryoid body formation for ESCs in 2iL or 2iLA at P6 and R-ESC (Reversing 2iLA at P3 to 2iL at P3). **f-i,** qRT-PCR for the formative genes *Dnmt3b*, *Dnmt31*, *Lef1* (**f**), and the telomere maintenance gene *Zscan4*. (**i**) or FACS for the differentiation marker T positive cells (**g**) or tetraploid complementation (**h**) for ESCs cultured in 2iL,2iLA, iErk or iFgfr1-3. **P<0.01, ***P<0.001, ***P<0.0001, student t-test. **(j)** Schematic summary of the pluripotency transition or genomic stability between naive and formative states. 2i-ESCs maintain naive state with genome instability, while supplement AlbuMAX (AX) in 2iL pushes naive to formative state transition with genomic stability. This lipid induced pluripotency transition is reversible by withdrawing AX from the 2iLA or by adding an additional p-Erk specific inhibitor or Fgfr inhibitor in 2iLA.

### Lipid-induced pluripotency transition is restrained through manipulation of Fgf/Erk signaling

We next asked whether the lipid-induced pluripotency transition can be restrained through inhibition of p-Erk1/2 using specific Erk inhibitors to counteract the effects of lipids (**Extended data Fig.8a**). The small molecule SCH772984 is a unique, selective and ATP competitive inhibitor of Erk1/2 signaling that acts by inhibiting phosphorylation of the Erk1/2 substrate p90 ribosomal S6 kinase [53]. SCH772984 was added to ESCs previously maintained in 2iLA medium (designated 2iLA+iErk) for 3 passages. ESCs in 2iLA+iErk medium displayed typical naïve colony morphology (**Fig. 6a**) and the proliferation rate slowed (**Extended data Fig.8c)**. Analyses of pluripotency marker genes confirmed decreased levels of formative genes including *Lef1* and *Dnmt3b/l* (**Fig. 6f**). These results demonstrate that lipid-induced formative pluripotency can be reversed to naïve state simply through manipulation of Erk1/2 signaling.

FGFR is upstream of the Erk1/2 signaling, and RNA-seq data confirmed increased expression levels for many FGF/MAPK/ERK pathway related genes in 2iLA conditions (**Extended data Fig.8a-8b**), including for *Fgf5* which is a validated epiblast or mEpiSC marker [54, 55]. Fgf5 interacts with membrane-bound receptor Fgfr1 to activate intracellular signaling cascades including PI3K-AKT, PLCγ, MAPK and STAT pathways [56]. Modulating FGF signaling was proposed to manipulate pluripotency state conversion [57], with inhibition of Fgfr as a means of maintaining murine ESCs in a more naïve state [8]. We added an inhibitor of Fgfr1/2/3, AZD4547 [58], to 2iLA medium (Designated 2iLA+iFgfr). Intriguingly, the AX-ESCs were reverted from formative to naïve state by this treatment (**Fig. 6a, 6f-6g**). The *in vivo* developmental potential of iErk/iFgfr-cultured ESCs was evaluated by tetraploid complementation. Both iErk and iFgfr cultured ESCs generated viable and healthy all-ESC pups with high efficiency (**Fig. 6h, Extended data Fig. 8d**), demonstrating that full pluripotency was preserved in these naïve ESCs that reverted from the formative state.

Supplementation of AX into 2iL medium significantly improves genomic stability and induced transition from naïve to formative pluripotency. To understand whether these phenotypes are intrinsically connected, genomic stability of ESCs cultured in 2iLA+iErk or 2iLA+iFgfr media was analyzed. Notably, ESCs cultured in 2iLA medium (formative pluripotency) maintained approximately 30% normal karyotypes, while ESCs cultured in either iErk or iFgfr conditions, even though appearing to be in a naïve pluripotent state, achieved over 40% and 20% normal karyotype cells at passage 12, respectively (**Extended data Fig. 8e**). We found that *Zscan4* expression levels, highly correlated with telomere length maintenance, were significantly higher in both iErk and iFgfr-ESCs compared to 2i-ESCs (**Fig. 6i**), indicating that telomere length is maintained in these ESCs. The results suggest that the improvement of genomic stability by lipid supplementation is independent of naïve or formative pluripotency state.

Therefore, formative AX-ESCs can fully revert to naïve state in terms of their colony morphology, transcript profile, and responsiveness to stimuli, by simply withdrawing AX from the culture medium. Addition of Fgf/Erk-specific inhibitors in 2iLA medium also reverts ESC cells to naïve state, retaining full pluripotency, but also with significant improvement of genomic stability (**Fig. 6j**), suggesting that lipid-induced pluripotency transition can be restrained through manipulating Fgf/Erk signaling.

### Male and female all-ESC mice are generated from de novo derived ESCs using AX-based medium

We next explored derivation of ESCs from E3.5-4.5 embryos from different strains using 2iLA and assessed the developmental potential by tetraploid complementation (**Fig. 7a**). After removal of the zona pellucida, E3.5-4.5 embryos were placed into a 96-well plate on feeders and cells expanded in 2iLA medium (**Extended data Fig. 8f**). Male and female ESC lines were derived from F1 (129X B6) using 2iLA medium. All-ESC mice can be generated from both male and female cell lines to obtain live pups, with 24.7% and 18.5% efficiency, respectively (live pups/embryos transferred, **Fig. 7b, Extended data Fig. 8f**). Both male and female all-ESC pups appeared normal and survived to adulthood. We also derived ESCs from the inbred C57B6 strain using 2iLA medium and found all-ESC pups can be generated from these, although not as efficiently as with the F1 hybrid strain (**Fig. 7b**). The outbred ICR line has been considered a non-permissive strain to derive ESCs [43, 59]. Using 2iLA medium, we efficiently derived ICR ESC lines from E3.5-E4.5 blastocysts. Both male and female ICR ESC lines generated all-ESC pups via tetraploid complementation with an efficiency of 6.7% for male and 12.5% for female (**Fig. 7b, Extended data Fig. 8f**). Our studies demonstrated that supplementing AX into 2iL medium (2iLA) enabled derivation of both male and female ESC lines with full potential to generate all-ESC pups, suggesting that lipids maintain genomic stability and developmental potential during ESC derivation.

**Fig. 7.**
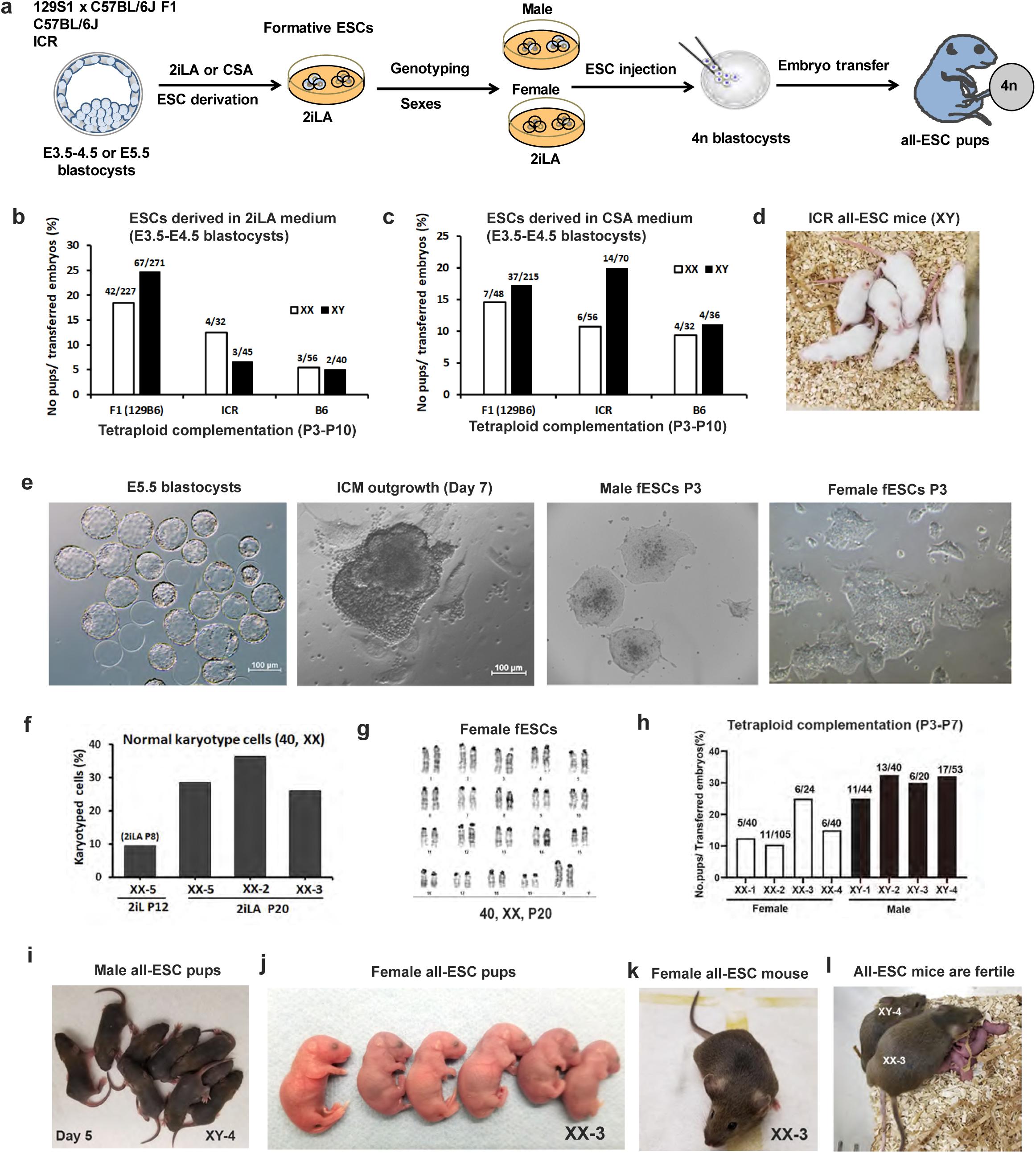
Generation of male and female all-ESC mice from de novo derived ESCs using AX-based medium. **a,** Schematic illustration of the experimental design. **b,** Tetraploid complementation assays for de novo derived ESCs (E3.5-E5.5 embryos) in AX-based medium. F1(129B6): hybrid F1 crossed with C57B6 females to 129S1 male; ICR: outbred strain; B6: inbred strain C57BL/6j; XX: female; XY: male. The numbers for each bar represent the No. pups/ No. embryos transferred. **c,** Tetraploid complementation assays for de novo derived ESCs (E3.5-E4.5 embryos) in CSA medium. **d,** A litter of all-ESC mice from a male ESC line derived from E4.5 ICR blastocysts in CSA medium. **e,** E5.5 blastocysts cultured in vitro (embryos hatching from the zona pellucida). The ICM outgrowth on feeders in 2iLA (second panel). The colonies of male fPSCs(third panel) or female fPSCs(fourth panel) derived from E5.5 blastocyst are shown in 2iLA at passage 3. Scale:100 μm.**f-g,** Karyotyping analyses of the embryo-derived female ESCs at P20. **h,** Tetraploid complementation assays for embryo-derived male and female ESCs. The numbers for each bar represent the No. pups/ No. embryos transferred. **i,** A litter of pups from XY-4 male cell line is shown at day 5 that was naturally delivered at term (9 pups/15 embryos transferred). **j,** A litter of embryo-derived female fPSCs (XX-3) is shown at day 1 (6 pups/24 embryos transferred). **k,** Adult female from the embryo-derived ESCs (XX-3). **l,** both male and female mice generated from the embryo-derived ESCs are fertile and produce normal litters. Bar scale in panel E: 100 μm.

Our results suggested that PD0325901(PD) is a major cause of genomic instability in cultured ESCs (**Fig. 4e**). To test if PD can be replaced by Erk1/2 inhibitors for ESC derivation, we substituted PD with the p-Erk1/2 inhibitor SCH772984 in 2iLA medium (CHIR/ SCH772984/LIF/AlbuMAX, designated CSA). ESC lines were readily generated from F1, ICR, and B6 strains with an efficiency as high as with 2iLA medium (**Extended data Fig. 8f**). The efficiency to obtain all-ESC pups were improved for the non-permissive strains of ICR (XX: 10.7%, XY: 20.0%) and B6 (XX: 9.4%, XY: 11.1%), while a similar efficiency as in 2iLA medium was achieved for the F1 strain (**Fig. 7c-d, Extended data Fig. 8g**). All-ESC mice generated from both male and female ESCs (F1, ICR, and B6) were normal and fertile (**Fig. 7c-7d**). Our results demonstrate that PD, the Mek1/2 inhibitor, can be substituted with Erk1/2 inhibitors for ESC derivation, suggesting that p-Erk1/2 direct inhibition and Wnt agonist are sufficient to preserve pluripotency for ESC derivation and maintenance in AX medium.

### Generation of all-ESC mice from ESCs de novo derived from late-stage blastocysts

Finally, we derived ESC lines directly from late-stage embryos (E5.5) using 2iLA medium and tested the cell lines for 4n competence through generation of all-ESC mice (**Fig. 7a**). We obtained and cultured 23 hatching blastocysts (E5.5) in the stage of embryonic escape from the zona pellucida (**Fig. 7e**). Embryos were placed individually into a 96-well plate on feeders and cells expanded in 2iLA medium. The ICM outgrowths emerged from all embryos at day 7 and cell lines were successfully established from each embryo (**Fig. 7e**). The colony morphology of these cell lines was distinct from ESCs cultured in 2iL medium and resembled the formative morphology of ESCs cultured in 2iLA medium (**Fig. 7e**). Notably, colonies from female cell lines were flatter and more irregular-shaped than those from male cell lines (**Fig. 7e**). The female cell lines derived in 2iLA medium were karyotyped and they maintained 25-35% normal 40, XX karyotypes at passage 20, while nearly all cells (over 90%) became aneuploid following 12 passage culture in 2iL medium, even if they were first cultured in 2iLA for 8 passages (**Fig. 7f, 7g**). Therefore, both male and female ESC lines can be efficiently derived from E5.5 blastocysts with improved genomic stability when maintained in 2iLA medium.

We next evaluated the developmental potential of male and female ESC lines derived from late stage blastocysts. All 4 male lines tested generated all-ESC pups with efficiencies from 25% to 30% (**Fig. 7h**). The pregnant surrogates carrying these embryos delivered naturally at term and the all-ESC pups were healthy and grew normally to fertile adults (**Fig. 7i**). Female ESC lines generated all-ESC pups through tetraploid complementation with efficiencies from 10% to 25% (**Fig. 7h**). Female all-ESC pups were normal and survived to adulthood (**Fig. 7j, 7k**), producing normal litters when crossed with male all-ESC mice (**Fig. 7l**). These results affirm that full pluripotency and 4n competence can be preserved in both male and female ESCs derived from late-stage blastocysts in 2iLA medium.

## DISCUSSION

Stabilization of pluripotency for both male and female ESCs during long-term culture would benefit research into the mechanisms of cell fate determination, epigenetic reprogramming, and modeling of early development using synthetic embryos. We show that supplementing 2i/LIF medium with lipid-rich albumin AlbuMAX significantly improves the genomic stability and developmental potential of murine ESCs. ESCs cultured in 2i/LIF+AX (2iLA) medium can be propagated in a state highly similar to formative pluripotency, which can be reversed to naïve pluripotency through withdrawing AX or adding additional FGF/Erk-specific inhibitors. Mechanistically, lipids directly stimulate Mek-mediated Erk2 phosphorylation, leading to exit of naïve state and establishing formative pluripotency for ESCs cultured in 2iLA medium. Lipid metabolism through β-oxidation is dispensable for transition from naïve to formative state. Lipid metabolism reduces the lipogenesis and amino acid biosynthesis and promotes non-canonical TCA metabolites recently reported to be involved in pluripotency transition [60]. Lipid metabolism promotes nucleotide and Acyl-CoA biosynthesis, enhances the expression of ZSCAN4 and DNMT3s that are involved in the maintenance of telomere length and DNA methylation, thereby improving genome stability during long-term culture. Stimulated Erk2 activity by lipids also alleviates X chromosome loss and possibly trisomy for ESCs cultured in 2iLA medium. The dual role of lipids on genome stability and pluripotency facilitates the preservation of 4n competency of murine ESCs for both sexes during long-term culture in vitro (**Extended data Fig. 9**). Successful generation of healthy, fertile female all-ESC mice from de novo derived ESCs demonstrates that AX-based culture media support derivation of fully potent murine ESCs of both sexes. Interestingly, in this AX-based system, PD, the Mek inhibitor, can be substituted with an Erk inhibitor (CSA), for efficient ESC derivation from various mouse strains.

It is known that lipids serve as signaling molecules in regulation of the Ras-Raf-Mek-Erk pathway [4, 61]. Our data show that Erk1/2 rapidly senses the addition of AX to elevate p-Erk1/2 independently of upstream Raf and Mek (**Fig. 5b**), and Mek1/2 is essential for the phosphorylation of Erk1/2 (**Extended data Fig.7h**), indicating that lipids regulating Erk2 phosphorylation might act through changing Mek1/2 kinase domain conformation, which is thought to be inhibited by PD. However, whether this is an effect induced by specific types of lipids needs to be further studied. While lipid-induced fPSCs share common features of formative pluripotency, they are distinct from other fPSCs reported in their transcriptional profiles, X chromosome inactivation, and developmental potential, suggesting that fPSCs can exist in sub-states reflecting the developmental continuum of pluripotency. We demonstrated that the lipid-induced fPSCs retain full potency to generate “all-ESC” mice, supporting the notion that fPSCs still possess unrestricted developmental potential.

The discovery of the 2i system [8] revolutionized both stem cell culture and derivation, not only for mice but also other mammalian species including humans [23-26, 62, 63]. Yet the 2i system has also been found to cause irreversible genetic and epigenetic changes in murine ESCs [28, 29]. The modified 2i systems (2iLA and CSA), supplemented with lipid-rich albumin, can efficiently prevent the detrimental effects of 2i and preserve pluripotency while maintaining genomic stability. This improvement provides a reliable culture system for targeting of ESCs and generating genetically modified mice, and also for modeling developmental processes, which depend on the genetic and phenotypic fidelity of ESCs during propagation. Female cells are typically sensitive to 2i resulting in loss of an X chromosome and DNA methylation; thus, female ESC lines derived using 2i have lost developmental potential [28, 29]. The modified 2iLA system can efficiently derive female cell lines that retain the potential to generate fertile all-ESC mice, suggesting that the genetic and epigenetic fidelity has been preserved. Therefore, the 2iLA system can be used for de novo derivation of pluripotent stem cell lines including even for non-permissive strains.

In summary, our findings underscore the importance of lipids in cell culture media for maintenance of genomic, epigenomic and phenotypic integrity, and discover novel mechanisms governing pluripotent cell transitions that are conserved in early mammalian development from mice to humans.

## Data availability

The all the RNA-seq and RRBS data have been uploaded to the NBCI BioProject. The accession number is uploaded to the Gene Expression Omnibus website. The accession number is PRJNA806916.

## Methods

### Mice and embryos

Animals were housed and cared for according to a protocol approved by the IACUC of Weill Cornell Medical College (Protocol number: 2014-0061). Wild-type ICR mice were purchased from Taconic Farms (Germantown, NY); the 129S1 and C57B6 mice were purchased from Jackson Laboratories. Females were super-ovulated at 6–8 weeks with 0.1 ml CARD HyperOva (Cosmo Bio Co., Cat. No. KYD-010-EX) and 5 IU hCG (Human chorionic gonadotrophin, Sigma-Aldrich) at intervals of 48 hours. The females were mated individually to males and checked for the presence of a vaginal plug the following morning. Plugged females were sacrificed at 1.5 days post hCG injection to collect 2-cell embryos. Embryos were flushed from the oviducts with advanced KSOM (Cat # MR-101-D, Millipore) and cultured in KSOM medium in an incubator under 5% CO2 at 37°C until blastocyst stage for ESC injection.

### Blastocyst injection and tetraploid complementation

ESCs were trypsinized, resuspended in ESC medium and kept on ice. A flat tip microinjection pipette was used for ESC injections. ESCs were collected at the end of the injection pipette and 10–15 cells were injected into each blastocyst. The injected blastocysts were kept in KSOM until embryo transfer. Typically, ten injected blastocysts were transferred into each uterine horn of 2.5 dpc pseudo-pregnant ICR females. Tetraploid embryos were generated using 2-cell stage embryos flushed from the oviducts. The 2-cell embryos were subjected to electrofusion to induce tetraploidy. Fused embryos were moved to new KSOM micro drops covered with mineral oil and cultured to blastocyst stage until ESC injection.

### 2iL medium and 2iLA medium

2iL medium contains 1:1 mixture of DMEM/F12 and Neurobasal media supplemented with N2 (Gibco™, 17502048) and B27 (Gibco™, 17504044), L-glutamine (Millipore, TMS-002-C), 2-mercaptoethanol (Millipore, ES-007-E), 1x penicillin/streptomycin (Millipore, TMS-AB-2C) and LIF (Sigma-Aldrich, ESG1107), 1 µM PD0325901 (Sigma or Stemolecule) and 3 µM CHIR99021 (Sigma). 2iLA medium was made from 2iL medium supplemented with 1% AlbuMax (W/V, Gibco™, 11020039).

### ESC derivation and culture

ESCs were derived from hybrid F1 fertilized embryos derived by crossing C57BL/6J females and 129S1 males. The ESC derivation medium contains KnockOut^TM^ Dulbecco’s modified Eagle’s medium (Gibco, 10829-018), 20% KnockOut^TM^ Serum Replacement (Gibco, 10828), 10^3^ IU recombinant mouse LIF (Millipore, ESG1107), 2mM L-glutamine (Millipore, TMS-002-C), 1x penicillin/streptomycin (Millipore, TMS-AB-2C), 1x non-essential amino acids (Millipore, TMS-001-C), 1x nucleosides for ES cells (Millipore, ES-008-D), 1x β-mercaptoethanol (Millipore, ES-007-E), 1µM PD98059 (Promega, V1191) and 3 µM CHIR99021 (Sigma). The medium used for ESCs derivation was 2iLA medium which is standard 2iL medium supplemented with 1% AlbuMAX (w/v). Briefly, E3.5-E5.5 blastocysts were collected from the plugged females treated with CARDova/hCG. The zona pellucida was removed by briefly exposing the blastocysts to Tyrode’s acidic solution (Millipore, MR-004). Blastocysts with dissolved zona pellucida were picked up with a mouth pipette and washed in KSOM medium for 2-3 times. Blastocysts were then placed individually on 96-well plate on feeder layers with the derivation medium. After 5-7 days’ culture, the ICM outgrowth originated from the blastocysts were trypsinized in 20 µL 0.025% trypsin solution (Millipore, SM-2004-C) for 5 min, and 200 µL culture medium was added to stop the reaction. Colony expansion of ESCs proceeded from 48-well plates to 6-well plates with feeder cells. Cell aliquots were cryopreserved using Cell Culture Freezing Medium (Millipore, ES-002-D) and stored in liquid nitrogen. ESCs were routinely expanded and maintained as following: Before passaging, the plates were treated with EmbryoMax™ 0.1% gelatin solution (Cat # ES-006-B, Millipore) for at least 10 min, and the ESC culture media were added to the wells after removal of gelatin solution. ESCs were then placed on the wells with proper cell density and usually cultured for 2 days before next passaging.

### Mutant ESC lines

The X^GFP^X^Tomato^ ESC line was a gift from Dr. Konrad Hochedlinger and the *Erk1/2* mutant ESC lines were obtained from Dr. Danny Reinberg with the permission from Dr. Sylvain Meloche. The Nanog-GFP cell line was a gift from Dr. Rudolf Jaenisch. The dual reporter ESC line (MERVL-tdTomato and ZSCAN4c-GFP) was a gift from Dr.Jianlong Wang. The *Mek1*^f/f^ *Mek2*^-/-^ iPSC cell line was a gift from Dr. Konrad Hochedlinger, the *Mek1*^-/^*^-^Mek2*^-/-^ iPSC cell line was generated by adding Cre Recombinase Adenovirus (Vectorbiolabs, Cat. No: 1710) into *Mek1*^f/f^ *Mek2*^-/-^ iPSCs. After transfection, the cells were sorted for GFP expression by flow cytometry and the sorted cells were expanded; single cells were picked and expanded for identifying the *Mek1/2* double knockout cell line. The *Cpt1a*^-/-^ ESC cell line was generated by CRISPR-Cas9 using Neon transfection system, the sgRNA sequencing is CACCACGATAAGCCAGCTGGAGG.

### Deproteinization of AlbuMAX

AlbuMAX deproteinization was performed as described in Garcia-Gonzalo and Izpisú a Belmonte [4]. Briefly, 0.5 mL of a 100X solution of Trypsin from bovine pancreas (Sigma) was added to 50 mL of a 10% w/v solution of AlbuMAX in base medium (AX-Trypsin solution) without supplement and incubated for 30 minutes at 37℃ in a water bath. Then, 0.5ml of a 100X solution of Soybean Trypsin inhibitor was added to the AX-Trypsin solution and incubated at room temperature for 30 minutes. The deproteinized AlbuMAX solution was sterile filtered and added to 2iL to a final concentration of 1% w/v. Fatty acid-free BSA (Sigma, A3803) was added to 2iL medium to a final concentration of 0.1% w/v. Chemically defined lipid concentrate (CDLC, Thermo Fisher Scientific, 11905031) was added at a concentration 2% v/v into 2iL media.

### Karyotype analysis

Briefly, cultures were treated with Colcemid at final concentration of 0.05 μg/m. Following 60-90 min incubation, cells were trypsinized according to standard procedures, washed twice in 1X PBS, incubated in 0.075M KCl for 10 minutes at 37°C and fixed in chilled methanol-acetic acid (3:1). The fixed cell suspension was then dropped onto slides, stained in 0.08 µg/ml DAPI in 2xSSC for 3 minutes and mounted in antifade solution (Vectashield, Vector Labs). The stained slides were scanned using a Nikon E800 epifluorescence microscope equipped with imaging and digital karyotyping system from Applied Spectral Imaging (Carlsbad, CA). For each sample a minimum of 20 metaphases were captured. All metaphases were fully karyotyped and analyzed for chromosomal instability. The experiments were performed at the MSKCC Molecular Cytogenetics Core Facility.

### RNA-sequencing

Total genomic DNA or RNA was prepared from cultured ESCs using the DNeasy Blood and tissue kit (QIAGEN) or RNeasy Mini Kit (QIAGEN), respectively, following the manufacturer’s instructions. The RNA concentration and integrity were measured using a NanoDrop 2000 (Thermo Fisher Scientific) and Agilent 2100 Bioanalyzer (Agilent), respectively. The integrity of RNA was indicated by the RNA integrity number (RIN). RNA samples with sufficient concentration and RIN greater than 8.0 were further prepared for cDNA library preparation using poly-A selection and unstranded library preparation using a Truseq library preparation kit (Illumina) according to manufacturer’s instructions. DNA library was then sequenced using a NovaSeq 6000 - S1 Flow Cell, with pair-end reads, 2×50 cycles at the Weill Cornell Genomics Core Facility. Reads from RNA-seq were aligned to mouse genome version mm10 using TopHat. Gene counts were obtained using featureCounts to sort bam files, and only unique-mapping reads were used. Genes without any expression counts in any sample were discarded. Differentially gene expression analysis was performed using DESeq2 (version 1.4.5) R package that normalizes gene count data to transcription per million (TPM), and then detects differentially expressed genes (DEG) between 2i-ESCs and AX-ESCs groups (FDR < 0.1).

### RRBS-sequencing

Reduced Representation Bisulfite Sequencing is a modification of the original RRBS protocol [64] and the in-house developed ERRBS method [65] for base-pair resolution methylation sequencing analysis based on the use of a restriction enzyme to enrich for CpG fragments. 150 ng of RNA-free genomic DNA per sample was used for RBBS sequencing at Weill Cornell Medicine Epigenomics Core. Concentration of double stranded DNA (dsDNA) was determined using Qubit Fluorometer, Perkin Elmer Labchip GX or agarose gel electrophoresis to determine molecular weight. 100 million (M) read per sample on a single end read flow cell with 100 sequencing cycles (SR100) was used for differential methylation analysis. FASTQ files were generated by bcl2fastq (V2.17) and filtered for pass filter reads based on Illumina’s chastity filter. Sequencing adapters were trimmed by FLEXBAR (V2.4) [66], genomic alignments using Bismark (V0.14.4) [67] and Bowtie2 (V2.2.5) [68] to reference mm10, and per base CpG methylation metrics were calculated with a custom PERL script. CpGs at a minimum threshold coverage of 5 reads were used for downstream analysis.

### Western blotting assays

Cells were lysed into 1× SDS loading buffer (50 mM Tris-HCl pH 6.8,5% β-mercaptoethanol, 2% SDS, 0.01% bromophenol blue, 10% glycerol) followed by sonication (Bioruptor, 2 × 30 s at high setting). Proteins were resolved on a 5–15% gradient Tris–glycine SDS–PAGE gel and semi-dry-transferred to nitrocellulose membranes. The following primary antibodies were used at the indicated dilutions: Zscan4 (Millipore, AB4340); Dnmt3a (Abcam, ab2850), Dnmt3b (Abcam, ab2851), GAPDH (CST, 5174, 1:10,000); Erk1/2 (CST, 4695); p-Erk1/2 (CST, 9101); Nanog (CST, D2A3 XP®). Horseradish peroxidase (HRP)-conjugated secondary antibodies and the ECL prime western blotting system (GE Healthcare, RPN2232) were used for detection of primary antibodies. For the Erk1/2 &p-Erk1/2 or Mek1/2&p-Mek1/2, the equivalent samples were loaded on two parallel gels synchronously and then the gels were transferred to nitrocellulose membranes synchronously. Chemiluminescent signals were captured with a digital camera (Kindle Biosciences).

### Quantitative real-time PCR

ESCs were homogenized in RNA lysis buffer (RLT buffer, Qiagen) with 1% β-mercaptoethanol. RNA was then prepared following the RNeasy protocol (QIAGEN). DNA was digested using DNase I (QIAGEN) during the RNA extraction processes. The RNA concentration was measured by NanoDrop (Thermo Fisher Scientific). RNA (200 ng) was used for cDNA conversion using qScript Super Mix (Quanta Biosciences). After cDNA dilution with double-distilled H_2_O at a 1:10 ratio, the qPCR reaction was prepared by mixing the gene-specific primers and PowerUp SYBR Green Master Mix (Thermo Fisher Scientific;cat. no. A25778). qPCR was run using a QuantStudio 3 machine (Thermo Fisher Scientific).

### Flow cytometry

The attached cells were washed with 1x PBS and dissociated with Trypsin-EDTA (0.025%) at room temperature for 5 min. The same volume of Defined Trypsin Inhibitor (Gibco, R007100) was added to the detached cell suspension, incubated at room temperature for 5 min, and then washed with 1xPBS and centrifuged for the cell pellet. Samples without fluorescent reporter were used as the negative control. Finally, the cell pellet was suspended in MACS (1x PBS with 2mM EDTA and 0.5% BSA) with DAPI and applied to a flow cytometer for analysis (FACSymphony™ A5 or Fortessa, BD) at the Weill Cornell Medicine Flow Cytometry Core Facility.

### Telomere length measurement

The ScienCell Absolute Mouse Telomere Length Quantification qPCR Assay Kit (AMTLQ, Catalog #M8918) was used to directly measure the average telomere lengths. The telomere primer set recognizes and amplifies telomere sequences. The single copy reference (SCR) primer set recognizes and amplifies a 100 bp-long region on mouse chromosome 10 and serves as reference for data normalization. A genomic DNA sample with known telomere length served as a reference for calculating the telomere length of target samples.

### ESC directed differentiation

For mesendoderm differentiation, embryoid bodies (EBs) were formed in serum free differentiation media (SFD). SFD consisted of 75% IMDM (Gibco), 25% Ham’s F12 (Corning cellgro), 0.5X N2 (Gibco), 0.5X B27 (Gibco), 0.05% BSA (Gibco), 0.5mM Ascorbic Acid (Sigma), 2mM Glutamine (Corning), 0.45 mM monothioglycerol (Sigma), 100U/mL Penicillin and 0.1 mg/ml Streptomycin (Corning). On the first day of differentiation, cells were dissociated with Accutase (Sigma) and plated at 40,000 cells/ml in Petri dishes. Activin A (R&D Systems or Peprotech) was added at designed timepoints: day0 (0h), day1(24h), and day2 (48h) at a final concentration of 75 ng/ml. For Brachyury staining, EBs were dissociated with Accutase, fixed with 2% paraformaldehyde (Electron Microscopy Sciences) and stained with anti-Brachyury PE-conjugated Antibody (R&D) for 2h according to manufacturer instructions. After two washes, cells were analyzed using an Attune Nxt flow cytometer (Life Sciences).

### Liquid chromatography–mass spectrometry (LC-MS)

For all metabolite analyses, cells were seeded in 2iL or 2iLA media in 6-well plates. Metabolites were extracted with 1 ml ice-cold 80% methanol supplemented with 20 mM deuterated 2-hydroxyglutarate (D-2-hydroxyglutaric-2,3,3,4,4-d5 acid (d5-2HG)) as an internal standard. After overnight incubation at −80℃, lysates were harvested and centrifuged at 21,000g for 20 min to remove protein. Extracts were dried in an evaporator (GenevacEZ-2Elite) and resuspended by incubation at 30℃ for 2h in 50 ml of 40 mg ml^-1^ methoxyamine hydrochloride in pyridine. Metabolites were further derivatized by addition of 80 ml of MSTFA plus 1% TMCS (Thermo Scientific) and 70 ml ethyl acetate (Sigma) and incubated at 37℃ for 30 min. Samples were analyzed using an Agilent 7890A GC coupled to Agilent 5975C mass selective detector in the Metabolomics & Lipidomics facility at The Rockefeller University. The GC was operated in split less mode with constant helium gas flow at 1 ml/min. One microliter of derivatized metabolites was injected onto an HP-5MS column and the GC oven temperature ramped from 60℃ to 290℃ over 25 min. Peaks representing compounds of interest were extracted and integrated using MassHunter software (Agilent Technologies) and then normalized to both the internal standard (d5-2HG) peak area and the protein content of duplicate samples as determined by a BCA protein assay (Thermo Scientific).

## Acknowledgements

We thank Dr. Fuqian Geng for the help with western blots. We would like to thank the Epigenomics Core Facility and Genomic Resource Co-facility at Weill Cornell Medicine for RRBS-seq, ChIP-seq and RNA-seq, and the Metabolomics & Lipidomics facility at The Rockefeller University for Liquid chromatography–mass spectrometry. This work was supported by the New York State Stem Cell Science Program (NYSTEM; contract C32581GG to D.W.), grant no. R01 GM129380-01 and 1 R21 OD031973-01 from the National Institutes of Health (to D.W.).

## Author contributions

D.W., B.C. and T.E. conceived the project and D.W., L.Z., B.C. and T.E. wrote the paper. M.G. and L.Z. performed the in vitro differentiation experiments. D.W. performed the blastocyst injection and tetraploid complementation assay. Y.Q. and D.W. derived the ESC lines from blastocysts. L.Z. performed the in vitro cell-culture, FACS, real-time PCR, library preparation for RNA-seq, RRBS-seq and metabolic profiling. B.C. and A.S. performed the metabolic profiling, western blot, immunostaining, FACS, and real-time PCR. G.N. and L.Z. performed karyotyping. X.W. and Y.H. contributed to the bioinformatics analysis, D.J., R.K. and C.D.A. offered insightful discussions, helped interpret the results and edited the manuscript.

---

**Extended data Fig. 1.**
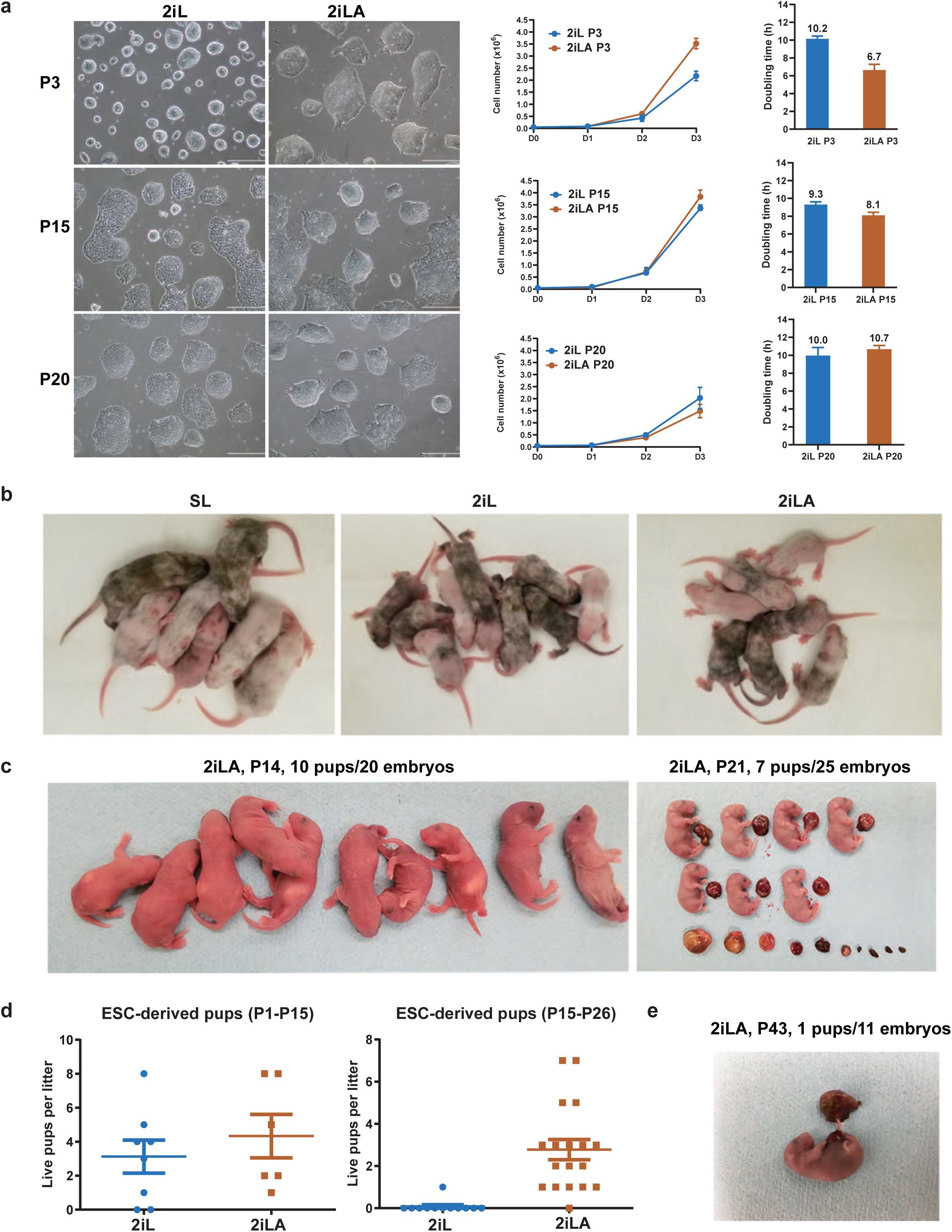
AX promotes ESC proliferation and developmental potency. **a,** Representative colonies, cell growth curves and doubling times of ESCs cultured in 2il or 2iLA at P3, 15, and P20. Scale bar in panel a: 25 µm. **b,** Representative chimeras derived from ESCs cultured in 2iL, 2iLA, or SL (serum+LIF) at P3. **c,** All-ESC pups from ESCs (ESC-1) cultured in 2iLA at P14 and P21. **d,** Dot plots quantifying the all-ESC pups obtained from ESCs (ESC-1) cultured in 2iL or 2iLA. **e,**one all-ESC pup obtained from the ESCs (ESC-1) cultured in 2iLA for 43 passages, which is almost three months in vitro.

**Extended data Fig. 2.**
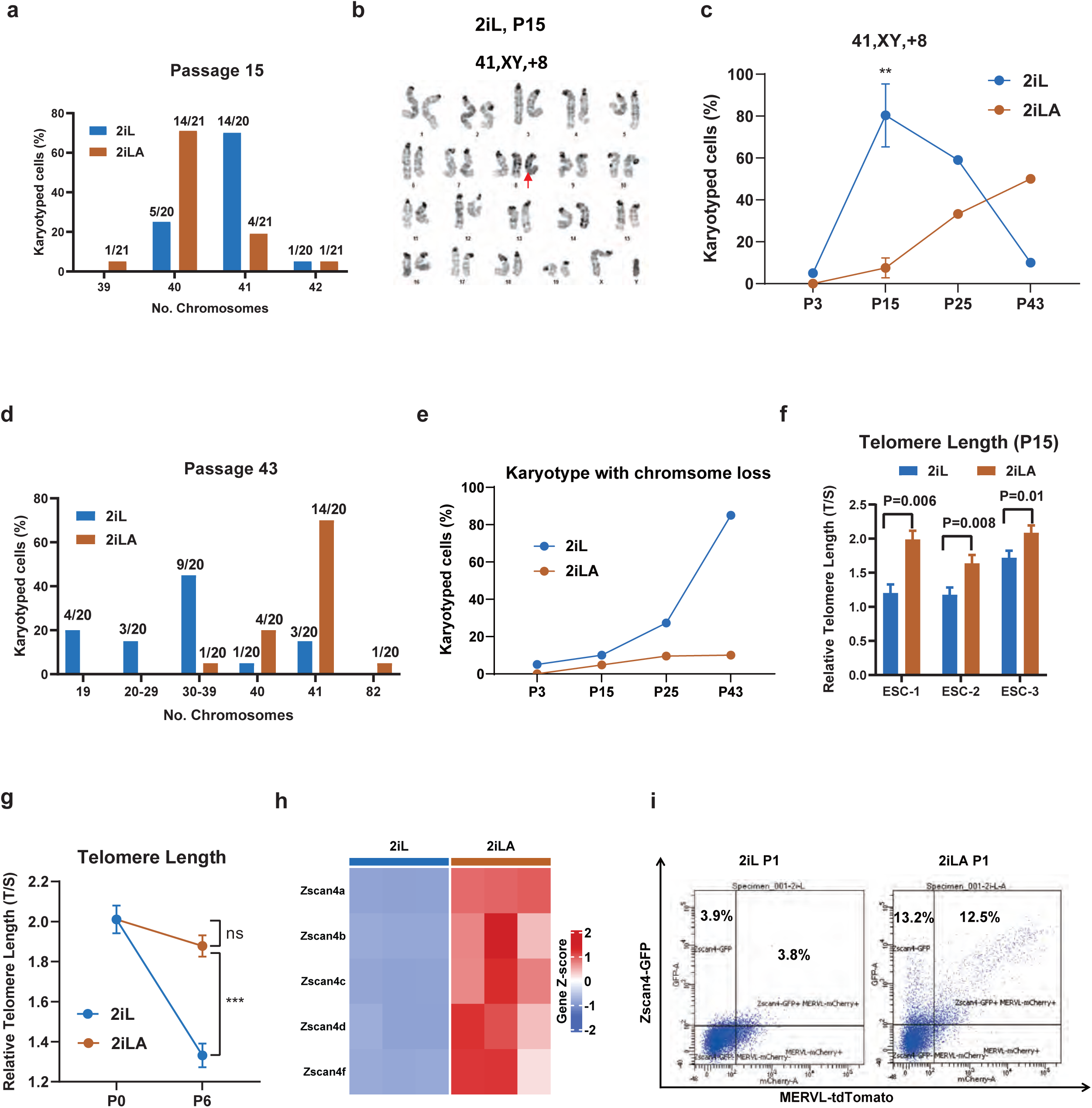
AX maintains genomic stability of ESCs. **a,** Karyotyping analyses for the number of chromosomes of the ESCs cultured in 2il and 2iLA at P15. **b,** Representative metaphase spread with trisomy 8 (41, XY, +8) from ESCs in 2iL at P15. Arrow: Trisomy 8. **c,** Karyotyping analysis for trisomy 8 in 2iL or 2iLA at different passages (n=∼20 metaphases at each passage for 2iL or 2iLA). **d,** Karyotyping analyses for the number of chromosomes of the ESCs cultured in 2il and 2iLA at P43. **e,** Analysis for karyotype with chromosome loss in 2iL and 2iLA at different passages (n=∼20 metaphases at each passage for 2iL or 2iLA). **f,** the telomere length analyzed by qPCR relative to a single copy gene(T/S) at passage 15 using three different ESC lines. Statistics were with Student’s t-test. **g,** Relative telomere length from P0 to P6, ***P<0.001, ns, non-significant, based on t-test. **h,** Heatmap of Zscan4 related transcripts from RNA-seq for ESCs cultured in 2iL or 2iLA at P3. **i,** FACS analysis to evaluate the Zscan4 positive cell percentage using the Zscan4-GFP and MERVL-tdTomato reporter cell line.

**Extended data Fig. 3.**
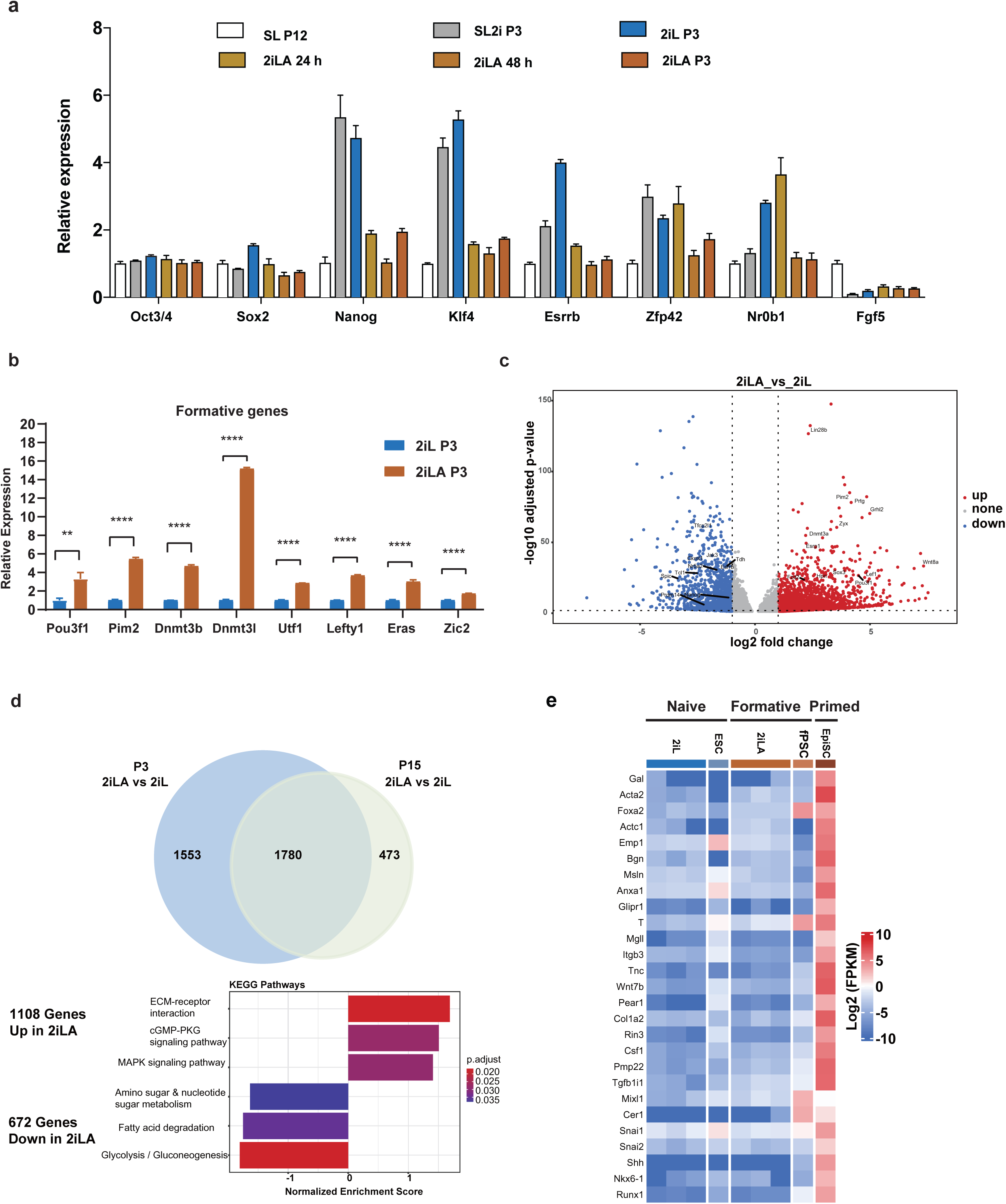
AX induces the transition of ESCs from naive to formative-like pluripotency. **a-b,** qRT-PCR for the naive or formative genes for ESCs cultured in 2iL, 2iLA, SL, or SL2i (a) or the formative genes in 2iL or 2iLA at P3 (b). **c,** Venn diagram plot of differentially expressed genes (DEGs) between 2iL and 2iLA ESCs at p3 or p15. Overlapping DEGs with FDR<0.1 & abs (FC)>1 between P3 and P15 and GO analysis for DEGs from bulk RNA-seq. n= three biological replicates for RNA-seq. **d,** Gene expression heatmap of primed genes, the mESC (Naive), fPSC (Formative), and EpiSC (Primed) are from published data [22].

**Extended data Fig. 4.**
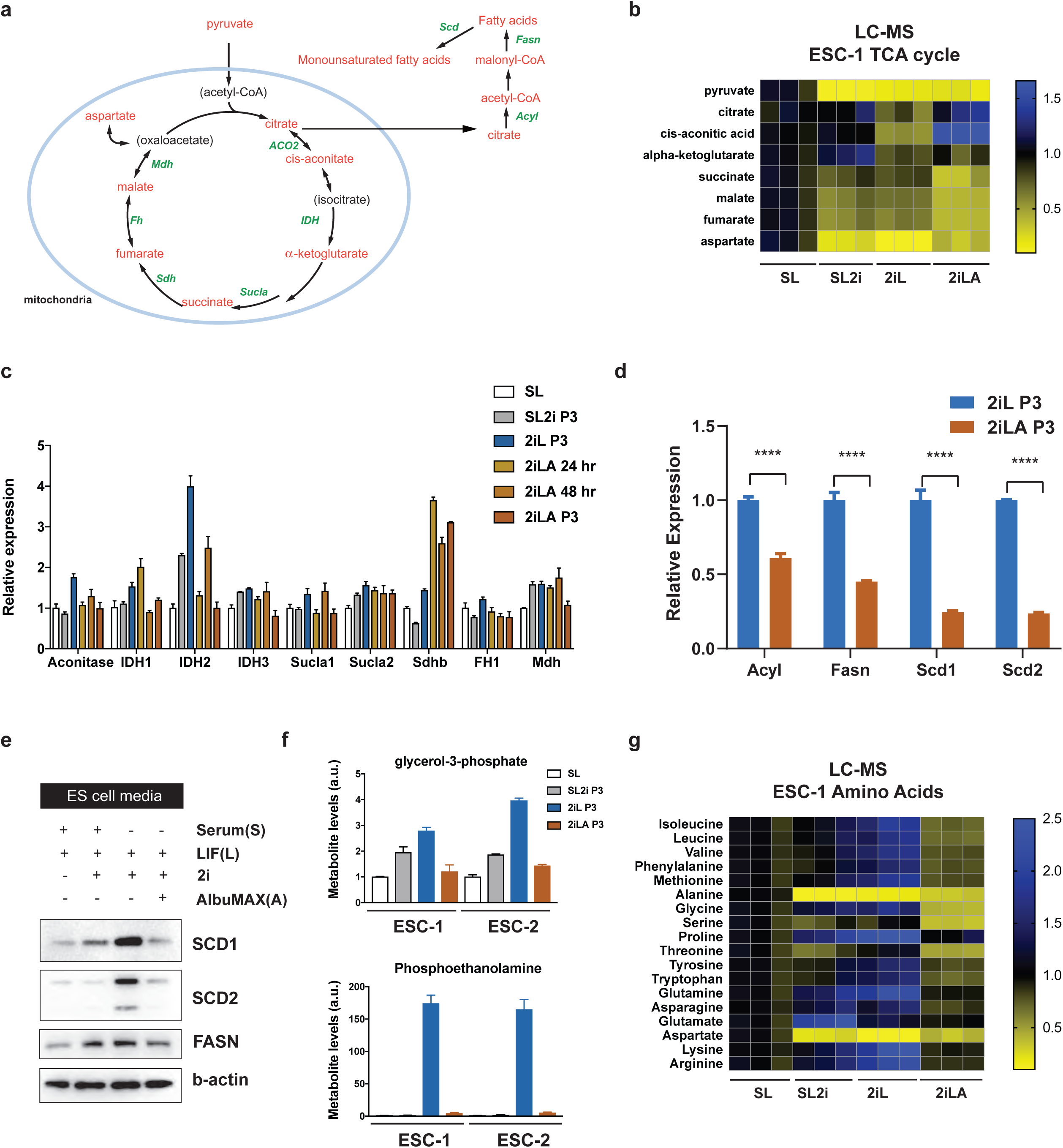
Lipids in AlbuMAX impact intracellular metabolism. **a,** Schematic of the tricarboxylic acid (TCA) cycle and de novo lipogenesis. **b,** LC-MS analysis of TCA cycle for ESCs cultured in SL, SL2i, 2iL or 2iLA. **c,** qRT-PCR for TCA related enzyme genes in SL, SL2i, 2iL, 2iLA 24hr, 2iLA 48hr, or 2iLA P3. **d-e,** Western blot (**d**) or qRT-PCR (**e**) for de novo lipogenesis related enzymes for ESCs in 2iL or 2iLA at P3. **f,** LC-MS analysis of the metabolites glycerol-3-phosphate or phosphoethanolamine for ESCs in SL, SL2i, 2iL or 2iLA. **g,** LC-MS analysis of amino acids for ESCs cultured in SL, SL2i, 2iL or 2iLA.

**Extended data Fig. 5.**
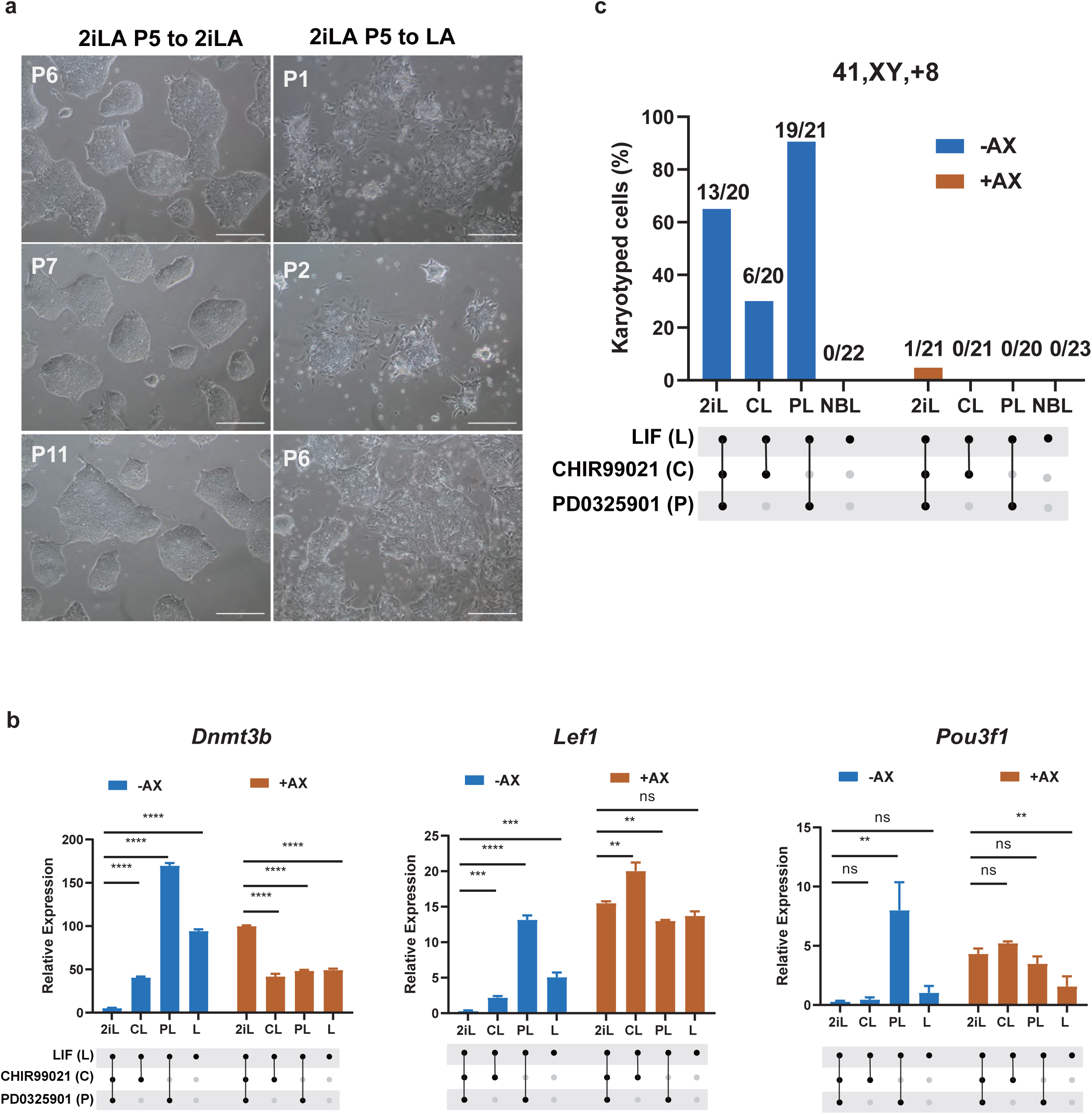
Formative-like pluripotency is maintained in concert with 2i. **a,** shown are representative colony morphologies of ESCs cultured in 2iLA (CHIR+PD+LIF+AlbuMAX) or LA (LIF+AlbuMax). Scale, 25 µm. **b,** Formative pluripotency marker genes *Dnmt3b*, *Lef1* and *Pou3f1* expression levels by qRT-PCR. **P<0.01, ***P<0.001, ****P<0.0001, ns: non-significant, based on t-test. **c,** Karyotyping analyses of trisomy 8 (41, XY, +8) for ESCs cultured in media with AlbuMAX (+AX) or without (-AX) after 15∼16 passages. Trisomy 8 karyotype/Total analysis metaphases. n=∼20 metaphases for each condition.

**Extended data Fig. 6.**
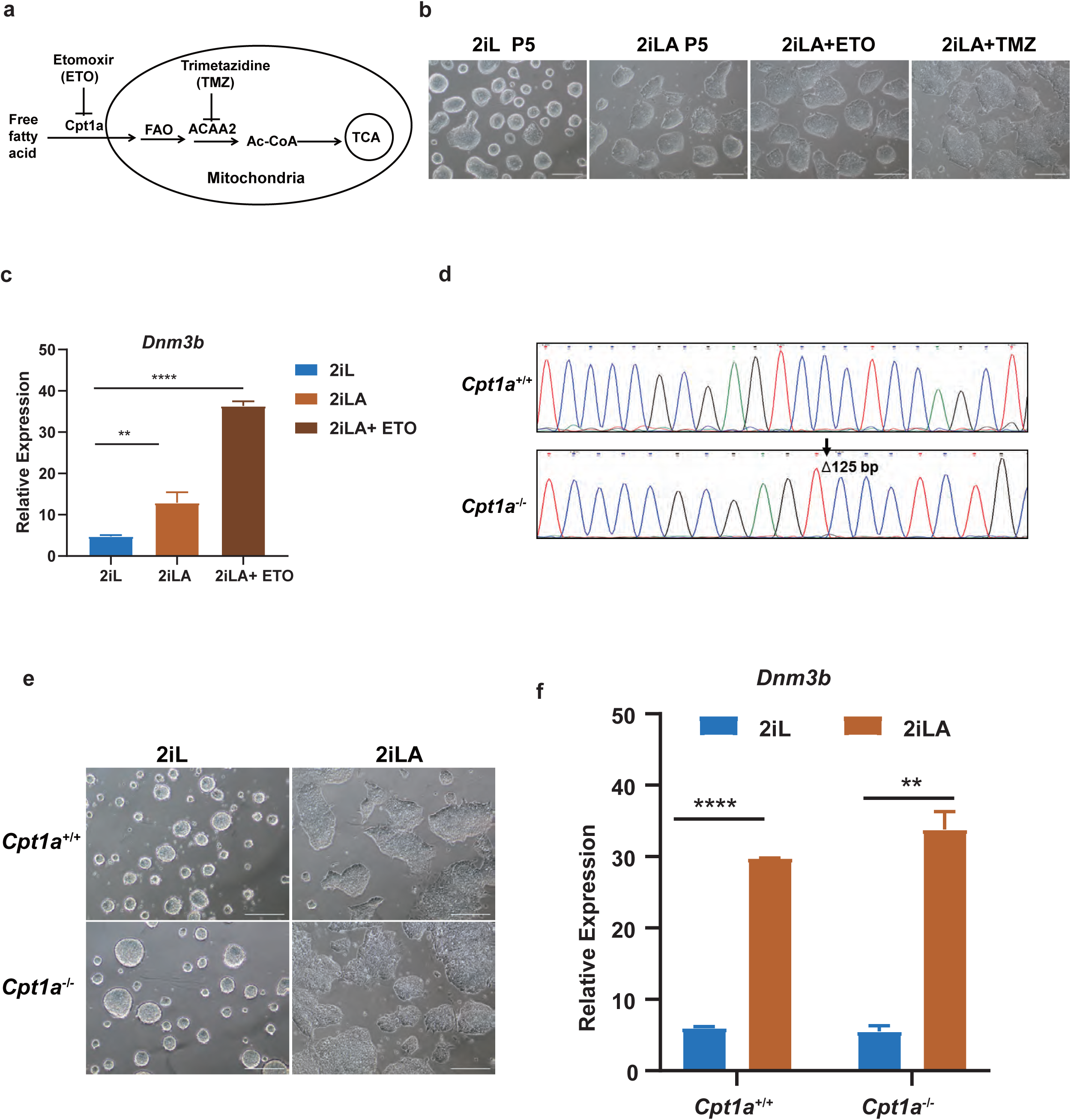
Fatty acid oxidation (FAO) is not required for lipid induced pluripotency transition. **a,** Schematic illustration of the experimental design for FAO pathway inhibition. **b,** Colony morphology after FAO pathway inhibition. Etomoxir (ETO, 100µM), Trimetazidine (TMZ, 200µM). **c,** Formative gene Dnmt3b expression after FAO pathway inhibition. Etomoxir (ETO, 100µM), Trimetazidine (TMZ, 200µM). **d,** Carnitine Palmitoyltransferase 1A (Cpt1a) deletion in mESCs, homologous deletion (the deletion region 125 bp spanning between exon3 and intron 3). **e,** Colony morphology after Cpt1a deletion. **f,** Formative gene *Dnmt3b* expression (qRT-PCR) after Cpt1a deletion.

**Extended data Fig. 7.**
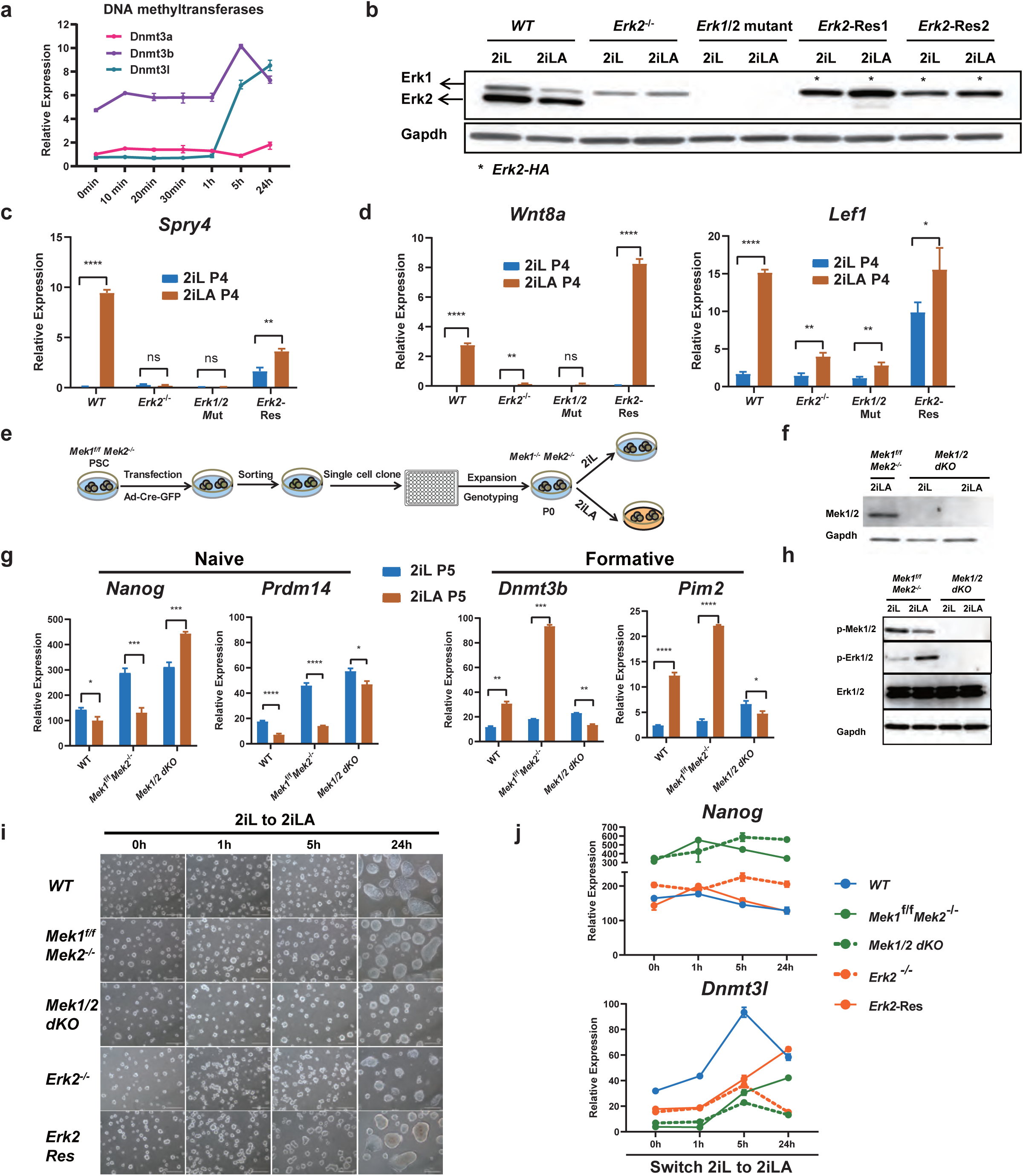
Mek/Erk signaling is essential for lipid-induced pluripotency transition. **a,** qRT-PCR for *Dnmts* transcript levels in ESCs after switching 2iL to 2iLA at different time points. **b,** Western blotting for Erk1/2 in *WT*, *Erk2^-/-^*, *Erk1/2* mutant, *Erk2-Res1* or *Erk2-Res2* ESCs in 2iL or 2iLA. Erk2-Res: Erk2-rescued in the *Erk1/2* mutant ESCs (Lentiviral-based shRNA against Erk1 in the *Erk2^-/-^* ESCs). **c-d,** qRT-PCR for the *Erk1/2* target gene *Spry4* (c), and the formative marker genes *Wnt8a* and *Lef1* for *WT*, *Erk2^-/-^*, *Erk1/2* mutant and *Erk2-Res* ESCs after 4 passages cultured in 2iL or 2iLA(d). **e-f,** Schematic illustration of the *Mek1/2* dKO ESCs generation and the experimental design (e), and Western blotting for validating the deletion Mek1/2 protein of the *Mek1/2* dKO ESCs cultured in 2iL or 2iLA (f). **g,** qRT-PCR for the naive genes *Nanog*, *Prdm14* and formative genes *Dnmt3b*, *Pim2* for *WT*, *Mek1^f/f^Mek2^-/-^*, *Mek1/2* dKO ESCs after 5 passages in 2iL or 2iLA. *P<0.05, **P<0.01, ***P<0.001, ****P<0.0001, t-test. **h,** Western blotting for the p-Mek1/2 or p-Erk1/2 protein level in *Mek1^-/-^Mek2^-/-^* dKO and its control PSCs. **i-j,** Colony morphology change (i) and qRT-PCR for naive gene *Nanog* and formative gene *Dnmt3l* (j) after swtiching 2iL to 2iLA at different time points for *WT*, M*ek1^f/f^Mek2^-/-^*, *Mek1/2* dKO, *Erk2^-/-^* or *Erk2-Res* ESCs.

**Extended data Fig. 8.**
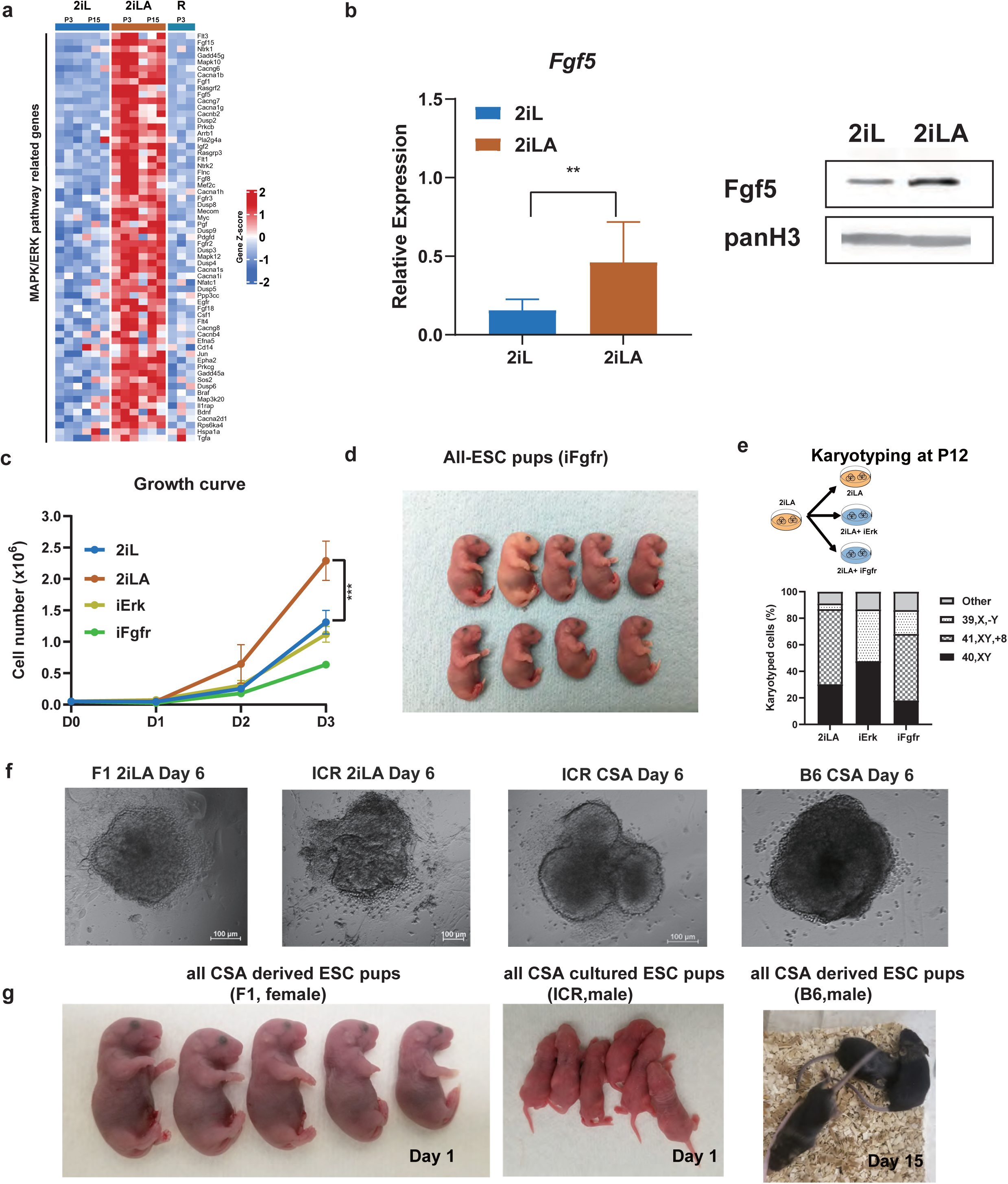
Fgf/Erk signaling manipulation in AX-based media and ESC derivation. **a,** heatmap showing transcript levels of the MAPK/ERK pathway related genes based on bulk RNA-seq for ESCs cultured in 2iL or 2iLA at P3. n=3 biological replicates. **b,** qRT-PCR or Western blotting assays (P3) of Fgf5 for ESC cultured in 2iL or 2iLA. **P<0.01, student t-test. **c,** the growth curve for the proliferation rates for ESCs cultured in 2iL, 2iLA, 2iLA+iErk, or 2iLA+iFgfr media. **d,** shown are nine pups derived from ESCs cultured in 2iLA+iFgfr1-3 from one surrogate recipient mouse (9 pups/ 20 embryos transferred) by tetraploid complementation. **e,** karyotyping analyses of ESCs cultured in 2iLA, 2iLA+iErk, or 2iLA+iFgfr media at P12. **f,** the ICM outgrowth from a F1 (129B6) or ICR E3.5 embryo on feeders at day 6 cultured in 2iLA or CSA. g, all-ESC pups generated for the CSA derived ESCs from F1 or B6 E3.5 blastocysts, and for CSA cultured ICR ESCs.

**Extended data Fig. 9.**
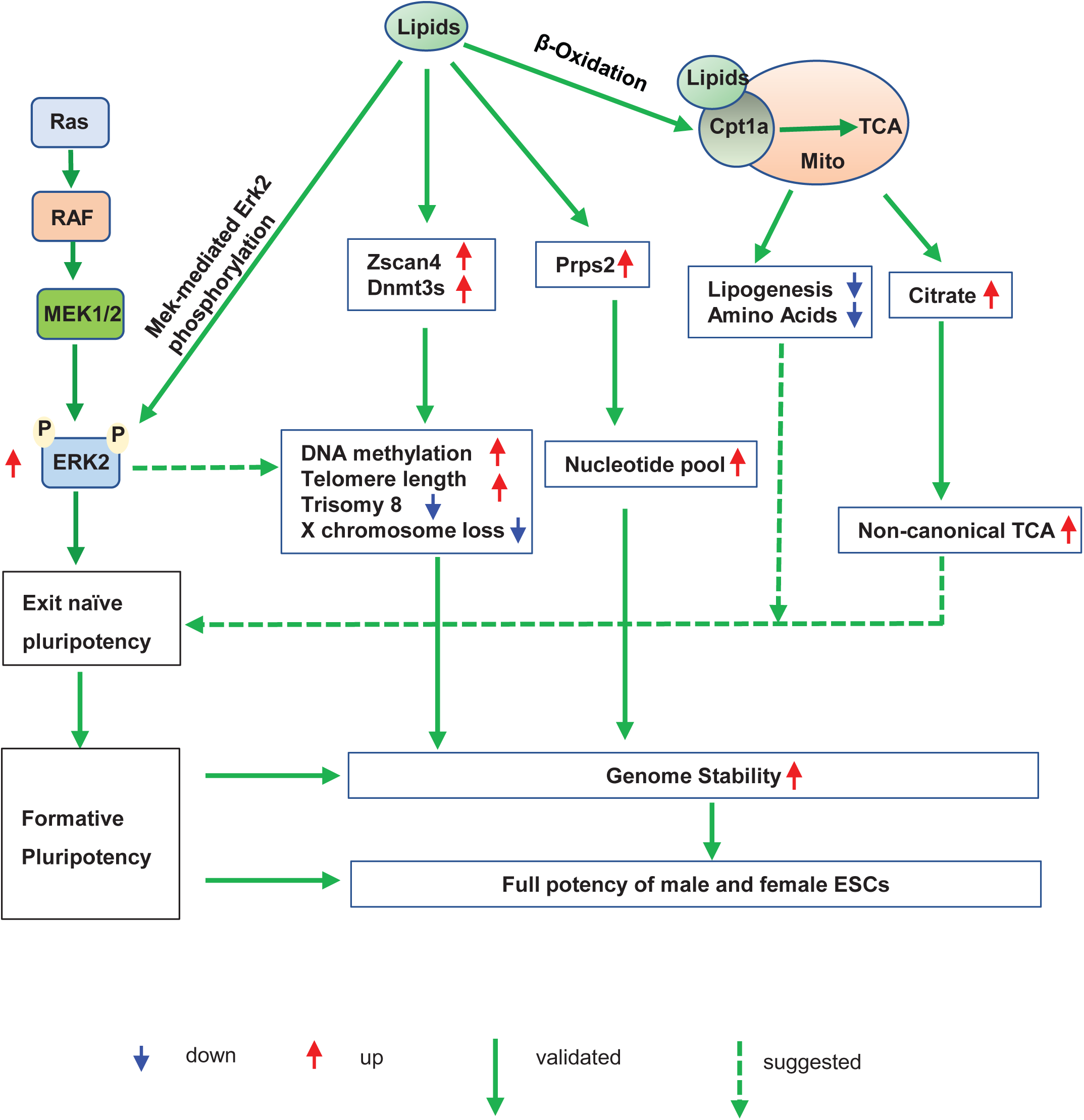
Schematic illustrating the dual role of lipids on genome stability and pluripotency of mouse embryonic stem cells. Lipids directly stimulate Mek-mediated Erk2 phosphorylation which leads to exit of naïve state and establishment of formative pluripotency for ESCs cultured in 2iLA medium. Lipid metabolism reduces the lipogenesis and amino acid biosynthesis and promotes non-canonical TCA metabolites associated with the exit of naïve pluripotency. However, lipid metabolism through β-oxidation is dispensable for transition from naïve to formative state. Lipids enhance the expression of ZSCAN4, DNMT3s and Prps2 that are involved in maintenance of telomere length, DNA methylation, and nucleotide pool, therefore promoting genome stability. Stimulation of Erk2 activity by lipids also alleviates X chromosome loss and trisomy 8 for ESCs cultured in 2iLA medium. The dual role of lipids on genome stability and pluripotency facilitates the preservation of full developmental potency of murine ESCs for both sexes during long-term culture in vitro.

